# The influence of molecular reach and diffusivity on the efficacy of membrane-confined reactions

**DOI:** 10.1101/439430

**Authors:** Ying Zhang, Lara Clemens, Jesse Goyette, Jun Allard, Omer Dushek, Samuel. A. Isaacson

**Affiliations:** Boston University, Department of Mathematics and Statistics, Boston, MA 02215; Center for Complex Biological Systems, University of California – Irvine, Irvine, CA; School of Medical Sciences, University of New South Wales, Sydney 2052, Australia; Sir William Dunn School of Pathology, University of Oxford, Oxford, OX1 3RE, U.K.

## Abstract

Signalling by surface receptors often relies on tethered reactions whereby an enzyme bound to the cytoplasmic tail of a receptor catalyses reactions on substrates within reach. The overall length and stiffness of the receptor tail, the enzyme, and the substrate determine a biophysical parameter termed the molecular reach of the reaction. This parameter determines the probability that the receptor-tethered-enzyme will contact the substrate, in the volume proximal to the membrane, when separated by different distances within the membrane plane. In this work we develop particle-based stochastic reaction-diffusion models to study the interplay between molecular reach and diffusion. We find that increasing the molecular reach can increase reaction efficacy for slowly diffusing receptors, while for rapidly diffusing receptors increasing molecular reach reduces reaction efficacy. In contrast, if reactions are forced to take place within the 2D plasma membrane instead of the 3D volume proximal to it, or if molecules diffuse in 3D, increasing molecular reach increases reaction efficacy for all diffusivities. We show results in the context of immune checkpoint receptors (PD-1 dephosphorylating CD28), a standard opposing kinase-phosphatase reaction, and a minimal two-particle model. The work highlights the importance of the 3D nature of many 2D membrane-confined interactions, illustrating a role for molecular reach in controlling biochemical reactions.

## INTRODUCTION

The ability of cells to sense their extracellular environment and make decisions relies on a diverse set of biochemical signalling reactions. Common to many of these reactions is the binding or tethering of an enzyme near its substrate before catalysis. Tethered signalling reactions are therefore controlled not only by binding affinities and catalytic specificities, but also by the properties of tethers which control the molecular reach of the reaction (defined below). Examples of tethered signalling reactions include those that take place on scaffolds [1, 2], and those that take place on the cytoplasmic tails of cell surface receptors. Tethering has also been used in synthetic biology to modulate endogenous signalling pathways [3, 4]. Although binding and catalytic reactions have been extensively studied experimentally and theoretically, the role of molecular reach is less well understood.

In the case of non-catalytic tyrosine-phosphorylated receptors (NTRs) [5], cytosolic enzymes first bind to their unstructured cytoplasmic tails before catalyzing reactions within reach. As a specific example, we consider the regulation of the NTR group member CD28 (Fig. 1A). This co-stimulatory receptor is expressed on T cells of the adaptive immune system, and is known to initiate signals important for their activation [6]. Phosphorylation of CD28 is mediated by the membrane-anchored SRC-family kinase LCK. It has been shown recently that CD28’s dephosphorylation is mediated by the NTR group member programmed cell death protein 1 (PD-1) [7]. This inhibitory receptor contains a tyrosine motif (ITSM) that serves as a docking site for the SH2 domain of the cytoplasmic tyrosine phosphatase SHP-2. When tethered to PD-1, SHP-2 is able to dephosphorylate tyrosines within reach, including those on CD28. Therefore, in addition to diffusion of these receptors within the membrane plane, it is expected that the tether will also play a role in controlling the ability of PD-1 to inhibit T cell activation.

**FIG 1.**
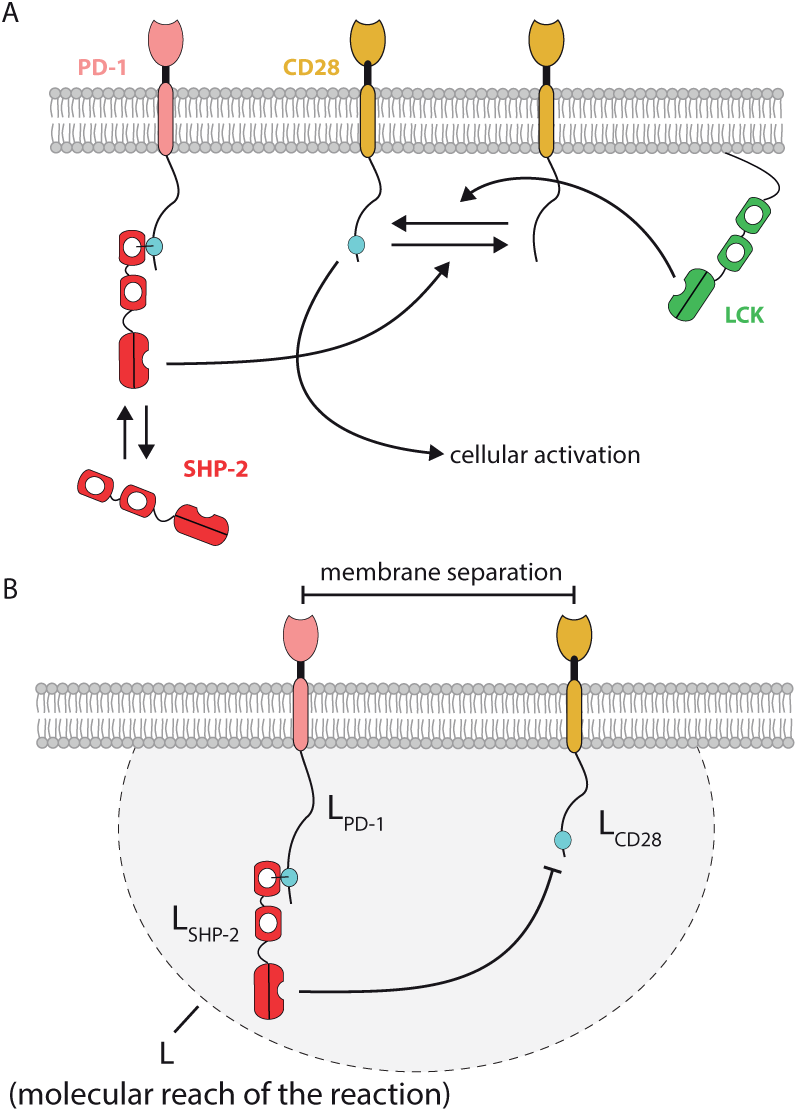
Illustration of tethered signalling reactions regulating the phosphorylation of the costimulatory surface receptor CD28 expressed on T cells. A) The membrane-anchored tyrosine kinase LCK is known to phosphorylate CD28. The cytoplasmic tyrosine phosphatase SHP-2 is known to dephosphorylate CD28 when tethered (or bound) to the cytoplasmic tail of the inhibitory receptor PD-1. B) The rate of CD28 dephosphorylation by SHP-2 will be controlled, in part, by the molecular reach of the reaction (*L*), with a larger reach generally increasing reaction rates when molecules are further apart. The molecular reach of the reaction will depend on the molecular reach of the individual components (*L*_PD-1_, *L*_SHP-2_, and *L*_CD28_). We estimate the molecular reach for this reaction to be *L ≈* 8.5 nm (see Methods).

In this example, the rate of CD28 dephosphorylation is expected to be influenced by the molecular reach of the reaction (Fig. 1B). Molecular reach determines the probability that the enzyme will contact the substrate when the two receptors are at a defined separation distance on the membrane. The overall molecular reach of the reaction is determined by the reach of the cytoplasmic tail of PD-1 (*L*_PD-1_), the reach of SHP-2 (*L*_SHP-2_), and the reach of the cytoplasmic tail of CD28 (*L*_CD28_). Here *L*_PD-1_, *L*_SHP-2_ and *L*_CD28_ will in turn depend on the respective length and stiffness properties of each component. By using the

worm-like-chain (WLC) polymer model, the overall molecular reach of the reaction can be defined as the square root of the squared sum of the individual reach parameters: *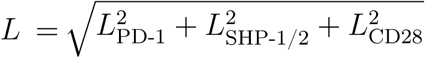* Experimental estimates of the molecular reach have yet to be reported, but we estimate the molecular reach for this reaction to be approximately *L* = 8.5 nm (see Methods for further details).

We note that SHP-2, and the homologous phosphatase SHP-1, are recruited to a variety of different receptors and act on a diverse set of substrates [8]. It follows that the molecular reach for SHP-2 (or SHP-1) catalysing a reaction on any given substrate (from different receptors) or from any given receptor (to different substrates) may exhibit wide variations. Indeed, the cytoplasmic tails of NTRs vary in their overall length [9].

To understand the role of molecular reach and diffusion in tethered signalling, we developed a particle-based convergent reaction-diffusion master equation (CRDME) model for the reaction and diffusion of individual receptors, kinases and phosphatases [10, 11]. Importantly, when simulating reactions between molecules confined to the 2D plasma membrane, we explicitly allowed their tails to explore the 3D volume proximal to the membrane by using a physiological 3D kernel that depends on the molecular reach (Fig. 1B).

Using our particle model, we first study the dephosphorylation of CD28 by PD-1 as the molecular reach of the reaction is varied. We find that the potency of PD-1 increases as the molecular reach increases for slowly diffusing receptors. In contrast, for rapidly diffusing receptors we find that increases in molecular reach reduce PD-1 potency. We show that this switch in potency as the molecular reach increases also holds in a commonly used biochemical model of reversible phosphorylation by kinases and phosphatases. In both biochemical models, we find that the switch is lost if membrane reactions are modelled using an idealised kernel that forces reactions within the 2D membrane plane. Using a simplified two-particle model that can be solved analytically, we reproduce these results. We then show that the switch arises from the constraint imposed by the molecules diffusing within the plasma membrane, which prevents the tethers from reaching all possible configurations where a reactive encounter could occur. Consistent with this, the switch is lost if molecules continue to interact using the 3D physiological kernel, but are instead allowed to diffuse in 3D. In this case, the region in which the molecules diffuse allows the tethers to sample all possible configurations in which a reactive encounter can occur. Our work highlights the 3D nature of 2D membrane-confined reactions, and suggests a possible unexpected role for molecular reach in controlling biochemical reactions.

## MATERIALS AND METHODS

### CRDME SSA Simulations

With the exception of our final simplified model, in which only one molecule diffuses, we study each of the biological models by Monte Carlo simulation of particle-based stochastic reaction-diffusion systems. Our simulation method is the convergent reaction-diffusion master equation (CRDME) stochastic simulation algorithm (SSA) [10, 11]. Here the diffusion of individual molecules is approximated by a continuous time random walk of the molecules hopping between voxels of a Cartesian mesh. First order reactions occur with an exponential clock, sampled independently for each possible first order reaction. Bimolecular reactions between two molecules occur with a separation dependent probability per time (derived from the Gaussian kernel *k*_cat_*σ*(*r*; *L*) for separation *r*, catalytic rate *k*_cat_, and molecular reach *L*, see the next section and [11]). In this way we approximate the diffusion and reactions of the molecules by a jump process. In SI Section S1 we provide a more detailed description of how the CRDME is formulated.

To study the first two models in the Results section, and the simplified two-particle model (Eq. S14), we used the CRDME SSA on a square/cubic domain with sides of length 300nm. The first two models had periodic boundary conditions on the sides of the square (cube), while the simplified two-particle model used a reflecting Neumann boundary condition, see Eq. S14. The domain was discretized into a Cartesian mesh of 2^16^ square voxels in 2D and 2^24^ cubic voxels in 3D. Each curve in Figs. 2B-D, Figs. 3C-H and Fig. S4 was estimated from 50000 simulations using the parameters in Tables I and II. For the first two models, simulations were run until individual trajectories reached steady-state. Our protocol for determining when steady-state was reached is described in SI Section S8. For the simplified model (Eq. S14), simulations were run until the two molecules reacted, with the corresponding reaction time then saved. As shown in SI S7, the qualitative dependence of the models on the diffusivity of molecules and the molecular reach of reactions was found to be relatively insensitive to the domain size (for molecular reaches much smaller than the domain width).

**FIG 2.**
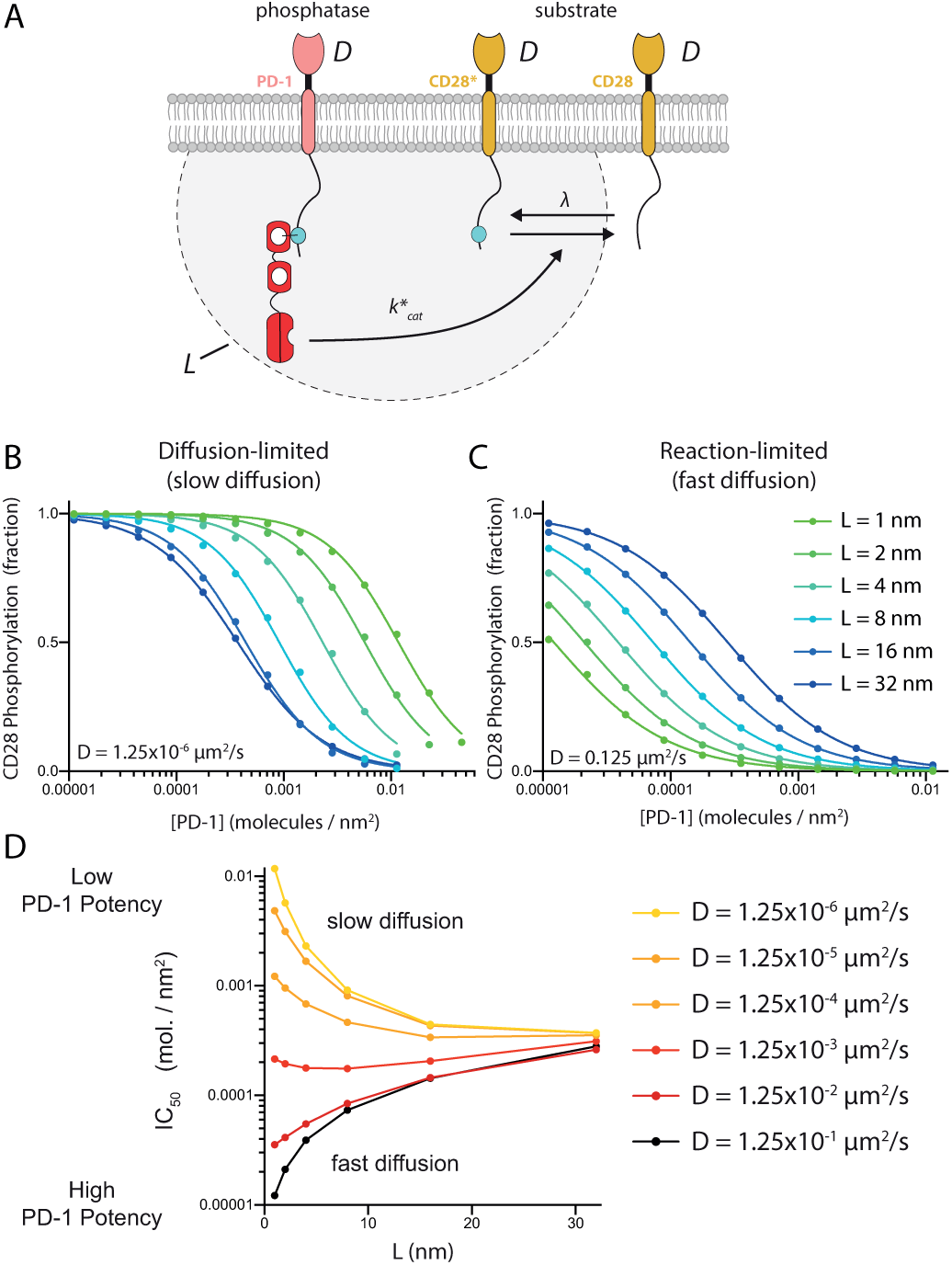
The potency of PD-1 receptor can increase or decrease as the molecular reach of the reaction increases depending on diffusivity. A) Schematic of species and biochemical reactions in our stochastic spatial model. B-C) Steady-state fraction of phosphorylated CD28 ([CD28*] / ([CD28*] + [CD28])) vs. [PD-1] for different values of the molecular reach for B) a smaller diffusion coefficient and C) a larger diffusion coefficient. D) Concentration of [PD-1] producing a 50% reduction in CD28 phosphorylation (also known as IC_50_) over the molecular reach of the reaction for different values of the diffusion coefficient. Parameters are summarised in Table I.

**FIG 3.**
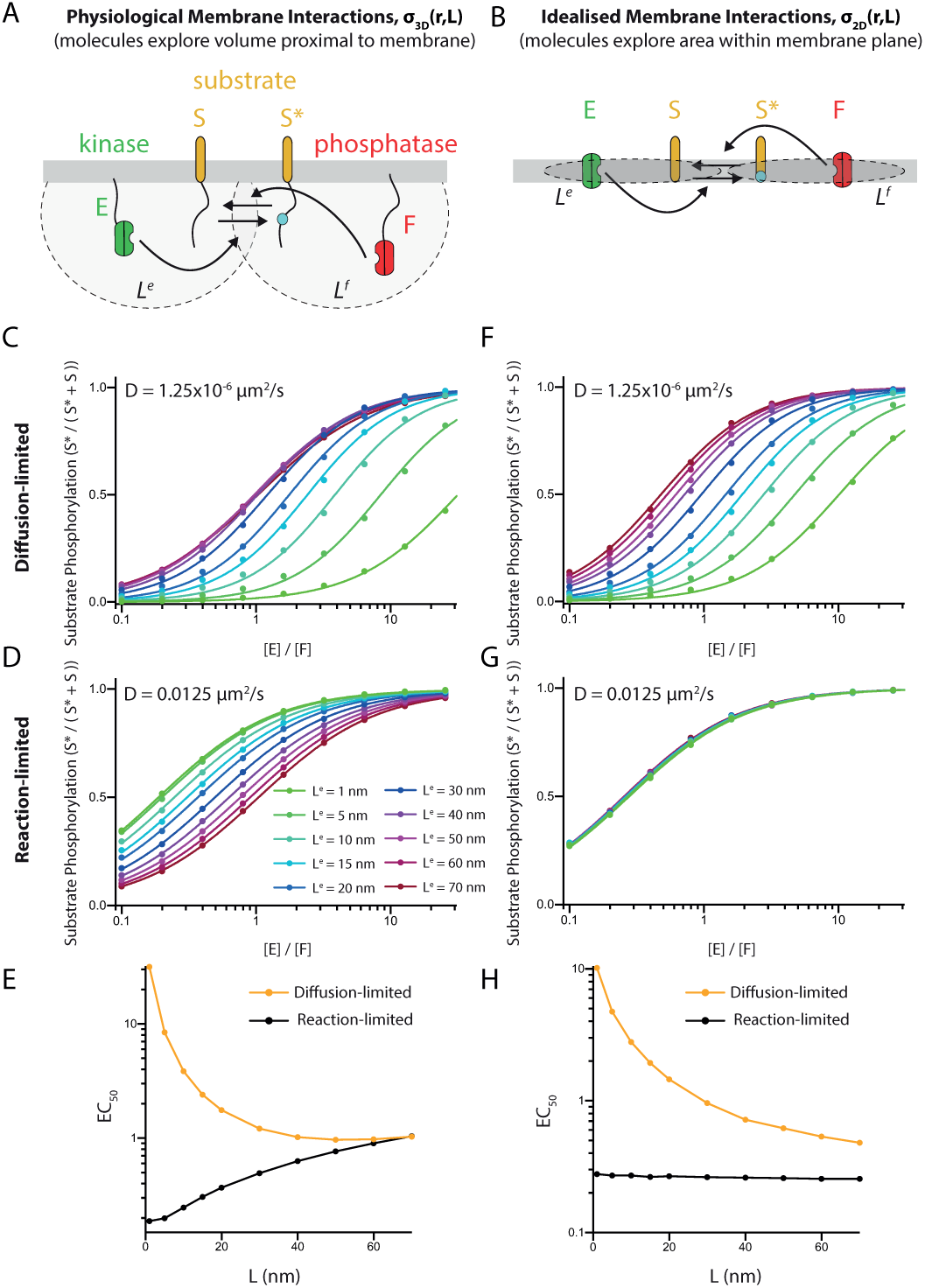
The switch in efficacy when increasing the molecular reach is only observed when explicitly allowing enzymes to explore the volume proximal to the membrane. A-B) Schematic of the bio-chemical model showing the reversible modification of a substrate by a kinase and phosphatase with reactions taking place A) within a volume proximal to the membrane or B) artificially confined to the plane of the membrane. The phosphorylation of the substrate is calculated in the steady-state for the physiological geometry (C,D) or the idealised geometry (F,G) when diffusion is limiting reactions (C,F) or when it is not limiting (D,G). Calculations are shown for different values of the molecular reach parameter for the kinase (legend in panel D applies to panels C,D,F,G). The potency of the kinase over the molecular reach is shown for the E) physiological and H) idealised geometry. All parameters are summarised in Table II. Note, when using the 2D kernel, *σ*_2D_, two-dimensional catalytic rates were obtained by rescaling the rates in Table II. If the square patch of membrane had side length *ℓ* we rescaled 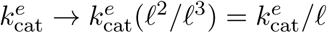 and 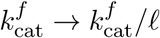.

**TABLE I.** Parameters for the PD-1 model (Fig. 2)

| Parameter | Description | Value |
| --- | --- | --- |
| $D$ | Diffusion coefficient | indicated $\mu\text{m}^2 \text{s}^{-1}$ |
| [PD-1] | PD-1 concentration | indicated $\text{nm}^{-2}$ |
| [CD28] | CD28 concentration | 0.0001 $\text{nm}^{-2}$ |
| $\lambda$ | Phosphorylation rate | 1.0 $\text{s}^{-1}$ |
| $k_{\text{cat}}^*$ | Catalytic efficiency | 0.1 $\mu\text{M}^{-1}\text{s}^{-1}$ |
| $L$ | Molecular reach | indicated nm |
| Domain | Periodic square | 300 nm × 300 nm |

**TABLE II.** Parameters for reversible phosphorylation model (Fig. 3)

| Parameter | Description | Value |
| --- | --- | --- |
| [S] | Substrate concentration | $100 \mu\text{m}^{-2}$ |
| [E] | Kinase concentration | indicated |
| [F] | Phosphatase concentration | $112 \mu\text{m}^{-2}$ |
| $D^s$ | Substrate diffusivity | indicated $\mu\text{m}^2 \text{s}^{-1}$ |
| $D^e$ | Kinase diffusivity | indicated $\mu\text{m}^2 \text{s}^{-1}$ |
| $D^f$ | Phosphatase diffusivity | $6.25 \times 10^{-4} \mu\text{m}^2 \text{s}^{-1}$ |
| $k_{\text{cat}}^e$ | Kinase catalytic efficiency | $0.04 \mu\text{M}^{-1} \text{s}^{-1}$ |
| $k_{\text{cat}}^f$ | Phosphatase catalytic efficiency | $0.01 \mu\text{M}^{-1} \text{s}^{-1}$ |
| $L^e$ | Kinase molecular reach | indicated nm |
| $L^f$ | Phosphatase molecular reach | 15 nm |
| Domain | Periodic square | $300 \text{ nm} \times 300 \text{ nm}$ |

### Derivation of probability density kernel (σ)

In our CRDME-based models, the kernel *σ*_3D_ determines the probability density that an individual tethered substrate (e.g. phosphorylation site on the cytoplasmic tail of CD28) will come in contact with an individual tethered enzyme (e.g. catalytic pocket of the phosphatase domain of SHP-2 tethered to the cytoplasmic tail of PD-1) at different separation distances between the membrane tether positions. That is, if the substrate’s tether is at position ***x*** in the membrane and the enzyme’s tether is a position ***y*** in the membrane, the separation distance between the tether positions is *r* = *|****x*** *−* ***y****|*. By assuming that the substrate and enzyme can be approximated by the worm-like-chain (WLC) polymer model, an analytical expression for the probability density kernel can be obtained [9, 12],

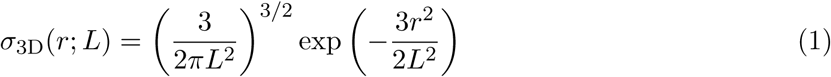

where *L* is the molecular reach for the reaction, and is given by the square root of the squared sum of the molecular reach of individual reaction components [9]. In the specific example of PD-1 dephosphorylating CD28 (Fig. 1-2), 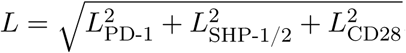 For the reversible phosphorylation model we consider in the Results (see Fig. 3),

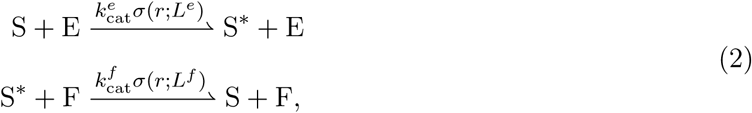

the molecular reach for the first (kinase) reaction would be *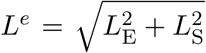*, and for the second (phosphatase) reaction *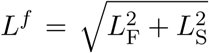*. Here *L*_E_ is the molecular reach of the kinase E, *L*_F_ is the molecular reach of the phosphatase F, and *L*_S_ is the molecular reach of the substrate S and phosphorylated substrate S*\**.

We note that the original derivation of Eq. 1 assumed that tethers explored free-space instead of the half-space imposed by the plasma membrane [9, 12]. Simulations revealed that the free-space kernel agrees with the half-space kernel up to a scaling constant (see SI Section S10) and therefore, to keep the models in the main text computationally efficient we have used Eq. 1.

In a number of simulations we replace *σ*_3D_ by an idealised two-dimensional interaction kernel to artificially force molecules to interact within the plane of the membrane, given by

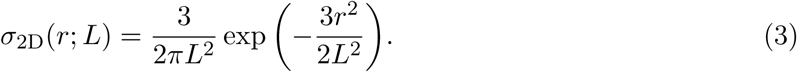

A plot of *σ*_3D_ and *σ*_2D_ is provided in Fig. S1.

### Estimating the molecular reach (L) for PD-1 dephosphorylating CD28

In the absence of experimental measurements, we approximate the molecular reach parameter for PD-1 dephosphorylating CD28. The molecular reach parameter for CD28 is simply *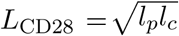* where *l*_*p*_ is the persistence length and *l*_*c*_ is the contour length. The persistence length for unstructured amino acid chains has previously been estimated to be *l*_*p*_ = 0.4 nm [12, 13]. The contour length of CD28 can be estimated using *l*_*c*_ =(0.4 nm) *×N*, where *N* is the number of amino acids between the membrane and the phosphorylated tyrosine, and 0.4 nm is the C_*α*_-C_*α*_ bond length in a polypeptide chain. The key activatory tyrosine in CD28 is the YMNM motif located 11 amino acids from the membrane and therefore *L*_CD28_ = 1.3 nm. Similarly, the molecular reach parameter for PD-1 is estimated to be *L*_PD-1_ = 3.0 nm, where the number of amino acids between the membrane and the tyrosine in the ITSM that binds SHP-2 is *N* = 56.

The molecular reach of the enzyme SHP-2 is more difficult to estimate because it is composed of 3 structured domains with flexible linkers: N-SH2(linker)C-SH2(linker)PTP (Fig. 1). Given that SHP-2 docks to its substrate primarily using the N-SH2 and catalyses reactions with its protein tyrosine phosphatase (PTP) catalytic domain, an upper bound for the molecular reach can be estimated by adding up the distances of the structured domains and the peptide linkers to obtain a maximum reach of 17.1 nm. However, the flexible linkers are unlikely to be maximally stretched and therefore a more realistic estimate is obtained by assuming a persistence length of 0.4 nm for the linkers that leads to an overall reach of 7.9 nm for SHP-2. We note that estimating the reach directly from crystal structure (PDB 2SHP) produces a value of 3.6 nm, but this value is for a single conformation of SHP-2.

In summary, the molecular reach of the reaction for SHP-2 bound PD-1 dephosphorylating CD28 can be approximated to be *L* = 8.5 nm.

## RESULTS

### A larger molecular reach can increase or decrease PD-1 receptor potency depending on diffusion

To investigate the influence of molecular reach on the ability of PD-1 to inhibit CD28, we developed a convergent reaction-diffusion master equation (CRDME) particle model (see Methods). The model included unphosphorylated CD28, phosphorylated CD28, and PD-1 bound to SHP-2, with all molecules able to diffuse in the plasma membrane (Fig. 2A). We explicitly included the effects of molecular reach by modelling the dephosphorylation of CD28 by PD-1 as a second order reaction whose rate was dependent on the separation distance between the molecules within the membrane (*r*),

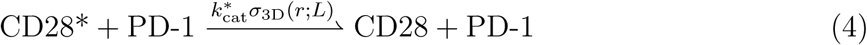

where *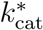* is the catalytic efficiency and *L* is the molecular reach of the reaction. The function *σ*_3D_ is the probability density (in units of molecules / nm^3^ or *µ*M) for finding the enzyme and substrate at the same location when their respective receptors are separated by a distance *r* within the plane of the plasma membrane. It depends only on the membrane-position of the receptors, but *accounts* for the diffusive motion of the tethered enzyme and substrate within the cytosol, see Methods. We calculate *σ*_3D_ under the assumption that PD-1 and CD28 can be approximated by the worm-like-chain polymer model, obtaining the Gaussian interaction given by Eq. 1 (see Methods). We focus on the effects of molecular reach for the dephosphorylation reactions and as such, we have introduced two simplifications to the model. First, we do not explicitly include the recruitment of SHP-2 to PD-1. Second, we do not explicitly model LCK molecules but instead model CD28 phosphorylation by a first order reaction 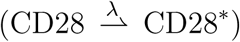 These simplifications are not expected to alter our conclusions, but decrease the computational complexity of the model by reducing the number of molecules that must be resolved in simulations.

As output of the model, we calculated the steady-state fraction of phosphorylated CD28 as the concentration of PD-1 was increased. We first focused on a situation where diffusion is minimal, which may be the case when immune receptors bind their ligands [14, 15], interact with the cytoskeleton [16, 17], and/or cluster [18]. As expected, increasing the concentration of PD-1 reduced phosphorylation of CD28 (Fig. 2B). In this case, we found that increasing the molecular reach of the reaction increased the potency of PD-1 so that fewer PD-1 molecules were necessary to achieve the same level of inhibition. Unexpectedly, when using a diffusion coefficient representative of free mobility on the plasma membrane for transmembrane receptors [16, 19], we found that increasing the molecular reach decreased the potency of PD-1 so that more PD-1 molecules were necessary to achieve the same level of inhibition (Fig. 2C).

We quantified the potency of PD-1 by calculating the concentration of PD-1 required to reduce the phosphorylation of CD28 by 50% (also known as IC_50_). A plot of IC_50_ over *L* shows that PD-1 potency increases for small but decreases for large diffusion coefficients, with a transition at intermediate values of the diffusion coefficient (*D* = 0.00125 *µ*m^2^/s) where potency is largely unchanged (Fig. 2D). Taken together, we find a switch in the effect of changing molecular reach, with larger reaches increasing receptor potency when diffusion is slow but decreasing receptor potency when diffusion is fast.

### Effect of molecular reach in physiological and idealised membrane reactions

A key novelty of our membrane bound protein reaction model is in accounting for reactions involving sites on molecular tails, which move through the volume proximal to the membrane. This is achieved through the use of the 3D interaction kernel *σ*_3D_, which accounts for the motion and stiffness properties of the tails, bound enzymes and substrates (see Methods). To determine the importance of the 3D kernel to the observed switch in reaction efficacy, we replaced the physiological kernel with an idealised 2D interaction kernel *σ*_2D_ (see Eq. 3).

This 2D kernel forced chemical interactions to only occur within the plane of the membrane (see Fig. S1), as in previous models [20]. To simulate this, and to generalise beyond the specific example of PD-1 acting on CD28, were-formulated the biochemistry of the model to a widely used scheme for the reversible modification of as ubstrate by a kinase and phosphatase [20–22] (Fig. 3A,B),

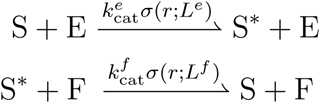

where S, E, and F are the substrate, kinase, and phosphatase, respectively, and ∗ indicates the phosphorylation modification (Fig. 3A-B). As before, we allowed for diffusion of all chemical species, and highlight that the rate of these enzymatic reactions is proportional to the catalytic efficacies (*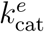* and *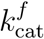*) multiplied by the probability densities (*σ*(*r*; *L*^*e*^) and *σ*(*r*; *L*^*f*^) for physiological 3D or idealised 2D interactions). The latter explicitly depends on the separation distance between the molecules in the simulation (*r*) and on the reaction molecular reach; *L*^*e*^ for the kinase phosphorylating the substrate and *L*^*f*^ for the phosphatase dephosphorylating the substrate.

We calculated the steady-state fraction of phosphorylated substrate as the number of kinase molecules was increased. Using the physiological 3D kernel, we reproduced the results for PD-1 (Fig. 2) where increasing the molecular reach increased the potency of the kinase when diffusion was slower but decreased its potency when diffusion was faster (Fig. 3C,D). When using the idealised 2D kernel, we found that increasing the molecular reach of the reaction increased the potency of the kinase when diffusion was slower (Fig. 3F) but when diffusion was faster it had no effect on the potency of the kinase (Fig. 3G). As before, we summarised these results by calculating the potency of the kinase as a function of the molecular reach for the physiological and idealised kernels (Fig. 3E,H). We confirmed that using the idealised 2D kernel in the PD-1 model of the last section also led the molecular reach to have a minimal effect in the reaction-limited, i.e. fast diffusion, regime (Fig. S2).

Taken together, these results highlight that the observed switching behavior in potency as molecular reach is increased is observed when using a physiological 3D kernel but not an idealised 2D kernel. We conclude that the 3D nature of 2D interactions can have profound effects on biochemical reaction rates.

### A minimal two-particle Doi model explains molecular reach phenotype

The preceding models demonstrate a clear switch in how the efficacy (quantified as potency) of tethered signalling reactions depends on molecular reach for large versus small diffusivities when molecules are confined to the 2D plasma membrane. They also suggest that such a switch may not be possible when molecules are forced to interact within the plane of the membrane. To understand what gives rise to this switch, and why it is not present when the molecules react in the membrane plane, we developed a simplified two-particle Doi model that could be solved analytically.

We consider a system containing just one A molecule and one B molecule, which can undergo the annihilation reaction

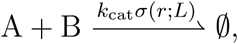

and assume that the A molecule is stationary and located at the origin, while the B molecule diffuses. We will consider three cases: the physiological model where the B molecule diffuses in 2D and tails interact in 3D (through the 3D Gaussian, *σ*_3D_(*r*; *L*)); a model where the B molecule diffuses in 2D but tails are forced to only interact in 2D (through the 2D Gaussian, *σ*_2D_(*r*; *L*)); and a model where the B molecule diffuses in 3D and tails interact in 3D (through the 3D Gaussian, *σ*_3D_(*r*; *L*)). In the remainder we denote these three combinations as the 2.5D, 2D, and 3D models respectively (Fig. 4).

**FIG 4.**
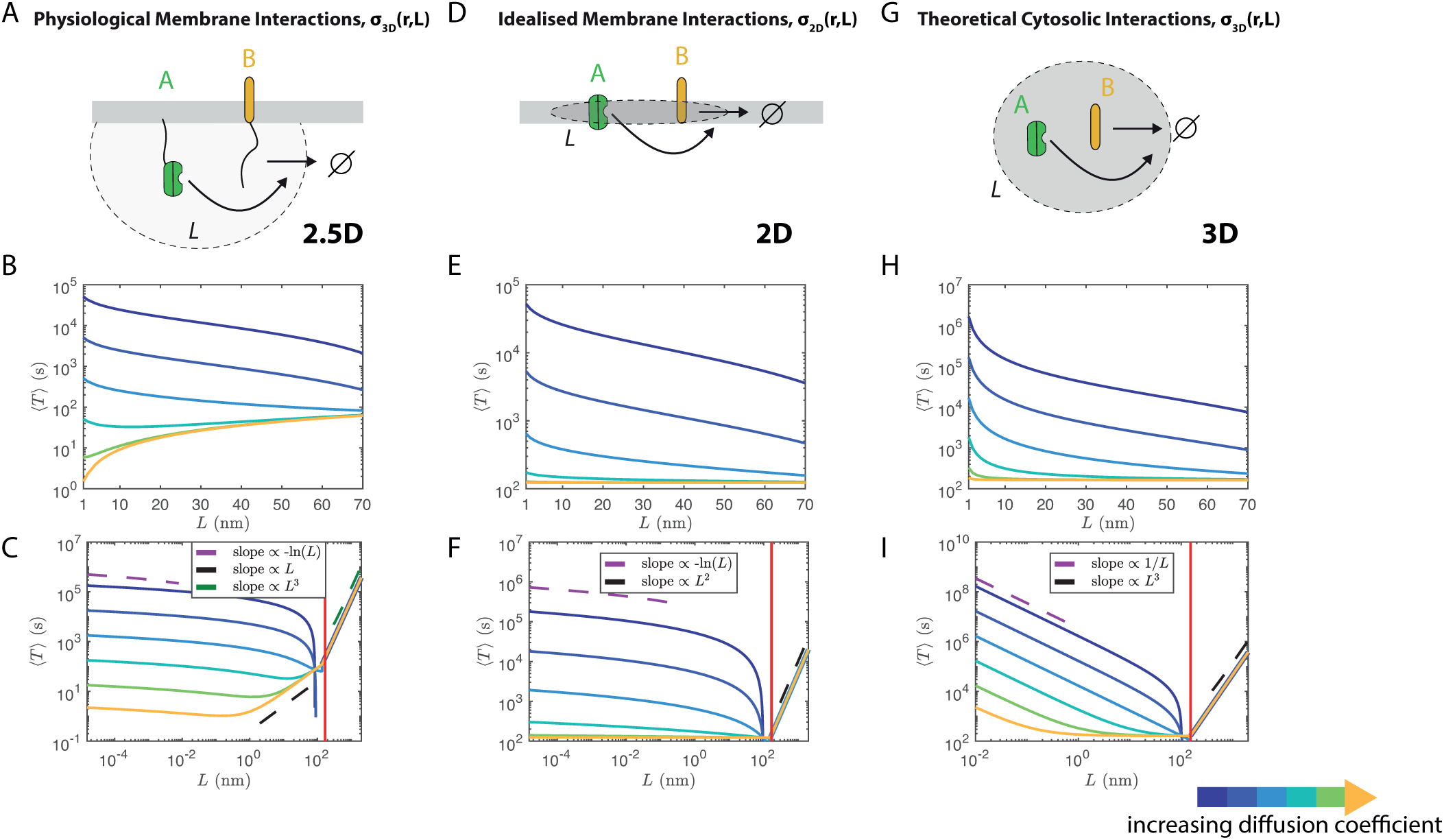
The well-mixed mean reaction time (MRT), ⟨*T*⟩, only demonstrates a switch in dependence on molecular reach for small vs. large diffusivities when considering membrane-bound molecules with cytosolic tails that react in 3D (2.5D model). In all figures solid lines correspond to the asymptotic expansions in Eq. 12a (B/C), Eq. 12b (E/F), or Eq. 12c (G/H). Dashed lines show general scaling behavior as a function of *L*. B) 2.5D model well-mixed MRT over physical parameter range. C) Same as B but showing an expanded range of *L* values. E) 2D model well-mixed MRT over physical parameter range. F) Same as E but showing an expanded range of *L* values. H) 3D model well-mixed MRT over physical parameter range. I) Same as H but showing an expanded range of *L* values. *In C, F and I an extreme range of molecular reach values is used to demonstrate the different scaling regimes of* ⟨*T* ⟩ *in L*. The vertical red solid line gives the *L* value such that *ε/R* = 1, corresponding to when the Doi interaction distance, *ε*, is equal to the domain radius, R. Note, as *ε* → *R* from below the asymptotic expansions break down since 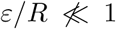. For B, C, H and I the catalytic rate *k*_cat_ is 0.1 *µ*M^*−*1^s^*−*1^. For E and F the catalytic rate *k*_cat_ is 0.1*πR*^2^*/*(4*πR*^3^*/*3) *µ*M^*−*1^s^*−*1^nm^*−*1^, the analogous 2D catalytic rate. Diffusion coefficients (blue to yellow): 1.25×10^*−*6^,1.25×10^*−*5^,1.25×10^*−*4^,1.25×10^*−*3^,1.25×10^*−*2^,0.1 *µ*m^2^/s

In the Doi model we assume the B molecule diffuses with diffusivity *D* within a circle (sphere) of radius *R* about the origin. *R* was chosen so that the area (volume) of the circle (sphere) was identical to that of the square (cube) with sides of length 300nm used in the preceding sections. We replace the Gaussian interaction *k*_cat_*σ*(*r*; *L*) by an approximating indicator function *λ* 𝟙_[0,*ε*]_(*r*), defined by

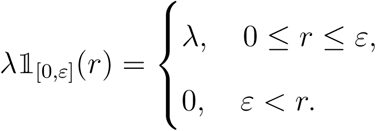

Here *λ* corresponds to the probability per time the molecules react when within a reaction-radius, *ε*, of each other.

The mean reaction time (MRT) *w*(*r*) for a diffusing molecule that is initially placed a distance *r* from the origin then satisfies

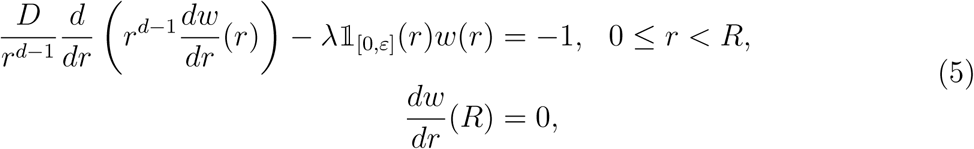

where *d* = 2 when the B molecule diffuses within a circular patch of membrane (2.5D and 2D models), and *d* = 3 when the B molecule diffuses within a spherical volume of cytosol (3D model). A no-flux boundary condition is used to prevent the B molecule from leaving the circle (sphere).

*λ* and *ε* are calculated by matching the total volume and the first moment of *σ*_3D_ for the 2.5D and 3D models. That is, given *k*_cat_ and *L*, we choose *λ* and *ε* such that

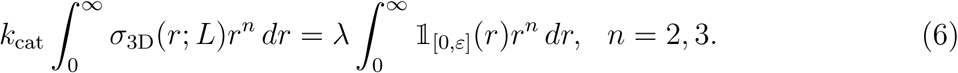

We find that

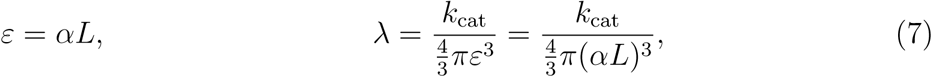

where 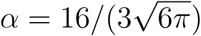.

When using *σ*_2D_ in the 2D model, *λ* and *ε* are calibrated by matching the total area and first moment, see Eqs. S21 and S22. In both calibrations we find that *ε* ∝ *L* (Eqs. 7 and S22), so that in the remainder we will interchangeably discuss changing *ε* or *L*.

We will focus on the well-mixed MRT, ⟨*T*⟩; the average time for the two molecules to react assuming the B molecule is initially placed randomly within the circle (sphere). It is given by

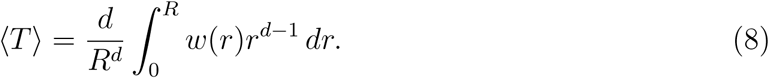

The 2.5D well-mixed MRT, ⟨*T*⟩, corresponding to substituting the solution of Eq. 5 into Eq. 8, is given by

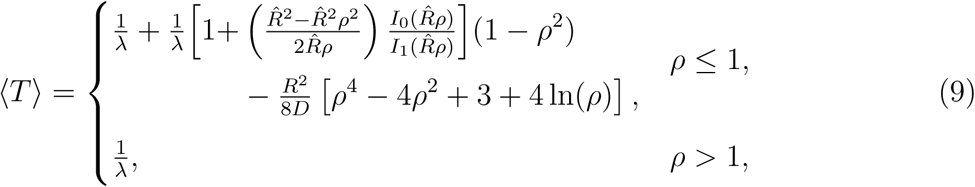

where ρ = ε/*R* and 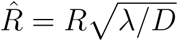.

To ensure that the replacement of the Gaussian interaction with the indicator function and immobility of the A molecule do not qualitatively change the behavior of the system, we compared (9) to a 2.5D CRDME model in which *both* molecules diffuse and react through *σ*_3D_. We demonstrate in SI Section S2 and SI Section S3 that ⟨*T*⟩ obtained from solutions of the Doi model (5) gives good qualitative agreement with the results of these CRDME SSA simulations.

To further simplify (9), we note that *L/R* is small in the biologically relevant parameter regime, so that *ρ* = *ε/R* = *αL/R* is also small. For *ρ* ≪ 1 we therefore expand Eq. 9 in *ρ* to obtain

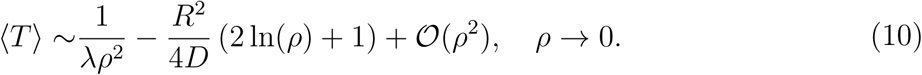

Using the calibrated parameters in Eq. 7, the 2.5D well-mixed MRT ⟨*T*⟩ can then be summarised by

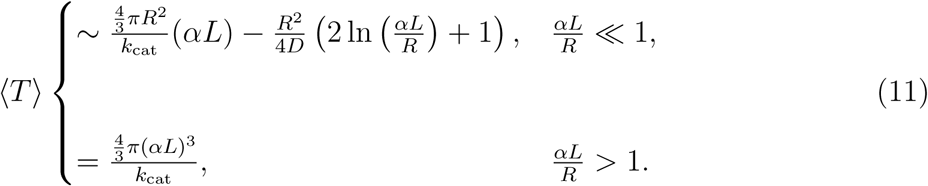

Using a similar approach to the preceding analysis (see SI Section S5 and SI Section S4), the well-mixed MRT of the Doi model (Eq. 5) can be found analytically for both of the 2D (Eq. 9 with calibration given by Eq. S22) and 3D (Eq. S17 with calibration given by Eq. 7) models. Their corresponding asymptotic expansions for *ρ* ≪ 1 are given by Eq. S23 and Eq. S20.

In summary, we find that over the physical range of molecular reach values, the exact solutions for ⟨*T*⟩ from the 2.5D, 2D, and 3D Doi models can be approximated by the asymptotic expansions

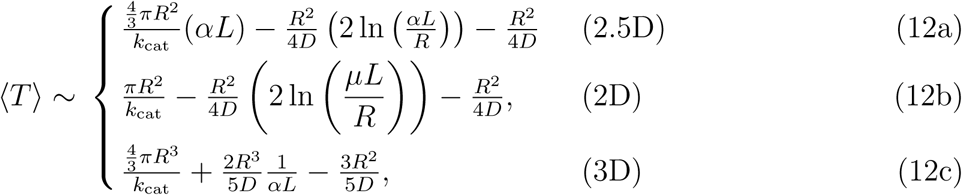

where 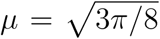. As shown in the SI, for physiological values of *L* and *D* these expansions agree well with numerical solutions to this model when using the original Gaussian interactions instead of the Doi indicator functions, see Fig. S6.

Fig. 4 plots the three asymptotic expansions as *L* and *D* are varied. Similar to our earlier models, in the physiological 2.5D case (Fig. 4B), we again see that when the diffusivity is small the reaction is most effective (⟨*T* ⟩ is smallest) for large values of the molecular reach, while for large diffusivities the reaction is most effective for small values of the molecular reach. In contrast, we observe that in both the 2D (Fig. 4E) and 3D (Fig. 4H) models increasing the reach always increases the reaction efficacy (decreases ⟨*T*⟩). We confirmed the latter result by simulating the biochemical model of the previous section in the fully 3D setting (molecules diffuse in 3D and interact using the 3D kernel) showing that, like the Doi model prediction, the potency of the kinase can only increase as the molecular reach increases (Fig. S3).

As we show in the SI Section S6, the first two terms in each of the three asymptotic expansions have a simple physical interpretation. We can write

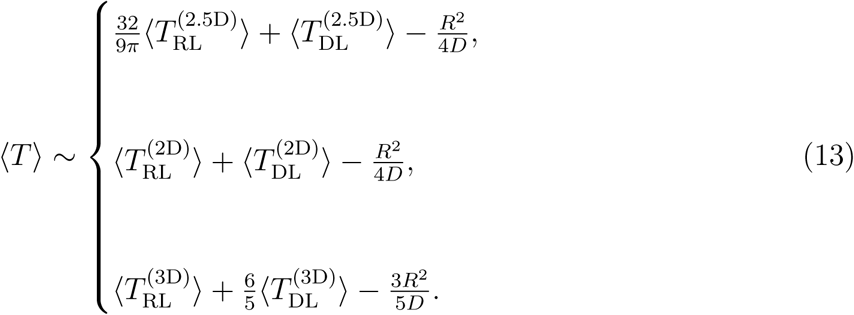

Here ⟨*T*_RL_⟩ denotes the reaction limited well-mixed MRT, corresponding to the well mixed MRT when diffusion is assumed to be infinitely fast, see Eq. S24. ⟨*T*_DL_⟩ denotes the leading order asymptotic expansion of the diffusion limited well-mixed MRT for *ε/R* ≪ 1, see Eq. S25. This corresponds to the diffusion limited regime, where the molecules are assumed to react instantly upon reaching a separation of *ε*. We therefore see that the well-mixed mean reaction time ⟨*T*⟩ can be (approximately) interpreted as the average time for the two molecules to get close enough to react (⟨*T*_DL_⟩), added to the average time for the two molecules to react when diffusion is sufficiently fast that the B molecule is always well-mixed (⟨*T*_RL_⟩).

The regime where ⟨*T*⟩ can increase as *L* increases only arises in the physiological 2.5D model. It is due to the reaction-limited well-mixed MRT, 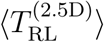, which is proportional to *L*. Eq. S24 shows that in both the 2D and 3D models the reaction limited well-mixed MRT is always independent of *L*, while Eq. S25 shows that the leading order diffusion limited well-mixed MRTs are decreasing in *L* for any diffusivity in all three models. The scaling of 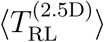 in *L* results from the use of a 3D Gaussian interaction (with units of inverse volume) in a planar region (with units of area), resulting in an effective well-mixed bimolecular reaction rate *k*_RL_ that scales like *L*^*−*1^. Since 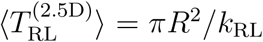, we find that 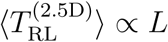 (see SI S6 for details).

We can interpret the (physical) differences between the 2.5D model and the 2D/3D models in the reaction-limited regime by considering the following points. For both the 2D and 3D models, changing the reach will change the shape of the Gaussian kernels, but it will not change the overall probability that the interaction sites are in contact (since in the reaction-limited regime the B molecule is still equally likely to be positioned at *any* of the possible locations that allow contact). In contrast, in the 2.5D model the position of the B molecule is restricted to a planar region despite the interaction occurring in the 3D volume proximal to the membrane. As the reach increases, an increasing fraction of the possible configurations where the interaction sites are in contact become inaccessible. The effective rate of the reaction will therefore decrease. In the opposite regime, where diffusion is rate limiting, increasing the reach will always increase reaction efficacy in each of the 2.5D, 2D and 3D models, as it becomes more likely that the interaction sites are in contact.

In summary, we find that for tethered signalling reactions the reaction time (i.e. ⟨*T*⟩) can exhibit a different functional dependence on molecular reach over physiological parameter regimes when diffusion is fast vs. slow. This arises from having 3D interactions between cytoplasmic tails of molecules confined to diffuse within a 2D membrane (2.5D model). We also find that when diffusion is sufficiently fast, the reaction time is independent of *L* for molecules diffusing and reacting in 3D (3D model), or diffusing and reacting purely in 2D (2D model). In contrast, the reaction time is still dependent on *L* for molecules diffusing in the membrane but reacting through the 3D interaction kernel (2.5D model). This illustrates how molecular reach in tethered signalling can reduce potency in 2D but not 3D geometries.

## DISCUSSION

Using a combination of spatial simulations and analytical calculations, we have examined the influence of molecular reach on membrane-confined reactions. Our key finding is that increases in molecular reach can increase reaction rates (or receptor potency) when diffusion is slow but decrease reaction rates (or receptor potency) when diffusion is fast. This switch is critically dependent on molecules diffusing in 2D but explicitly allowing them to react in the 3D volume proximal to the membrane using a 3D reaction kernel. The work underlines the importance of the 3D nature of 2D membrane-confined reactions.

### Reactions in 2D versus 3D

It is an open problem to understand how membrane confinement modulates receptor/ligand binding and biochemical reactions. Mathematical models of membrane reactions commonly restrict molecules to not only diffuse in 2D, but to react through 2D interactions [20, 23–26]. Although transmembrane domains (e.g. that localise PD-1 and CD28) and membrane-anchoring modifications (e.g. palmitoylation that localise LCK) restrict molecules to diffuse in the 2D membrane, their tethers allow them to explore a 3D cytoplasmic volume that is proximal to it. The switch in efficacy that we report critically relied on explicitly accounting for this through a physiological 3D kernel; using an idealised 2D kernel that forced molecules to interact within the plane of the membrane did not produce the switch.

### Modelling 3D reaction kernels for 2D membrane reactions

We have explored the molecular reach of the reaction primarily using a stationary Gaussian reaction kernel inspired by the worm-like-chain polymer model. It is likely that in some biological situations the polymer does not equilibrate quickly (stationary assumption) and/or the kernel is not Gaussian. We calculated that the stationary assumption is valid in our simulations (see SI Section S9) but this assumption will break down if, for example, longer tethers are simulated. A Gaussian kernel is expected to accurately capture the molecular reach of freely diffusing unstructured polypeptide chains such as the unstructured cytoplasmic tails of immune receptors [5]. However, there is evidence that the cytoplasmic tails of NTRs, including CD28, may have regulated interactions with the plasma membrane [27–29] which may lead to a non-Gaussian kernel. Similarly, a Gaussian kernel is expected to only be an approximation when applied to structured proteins like SHP-1/SHP-2 that contain multiple domains connected by flexible linkers. We note that experimental data of tethered dephosphorylation by SHP-1 was well-fit by a Gaussian kernel [9]. Nonetheless, careful consideration is needed when formulating a 3D reaction kernel and it may be feasible to determine the kernel using molecular dynamics or coarse-grained mesoscale simulations [30] that can be adapted to the specific molecules of interest.

### Implications to the biology of immune receptors

The ability of receptors within the NTR or immunoreceptor group [5] to regulate the phosphorylation of specific substrates is dependent on the signalling protein recruited by the receptor (e.g. SHP-2 in the case of PD-1), by the specificity of the signalling protein to the specific substrate (e.g. SHP-2 has the ability to dephosphorylate CD28 [7]), and the ability of the receptor and substrate to localise (e.g. PD-1/CD28 co-clustering [18]). In addition to these mechanisms, our work demonstrates that the molecular reach of a reaction may also control the ability of a receptor to regulate the phosphorylation state of the substrate and hence determine receptor potency. A key question that the present work raises is whether increasing the molecular reach will increase or decrease receptor potency? Although PD-1 and CD28 are expected to be mobile on resting T cells, their relative mobility within ligand-induced clusters has yet to be investigated. Our work indicates that increasing the molecular reach of this reaction will only increase PD-1 potency if their mobility is reduced within these ∼100 nm clusters [18].

### Experimental measurements

Tethered signalling depends on binding, catalysis, and on the molecular reach of the reaction. Although standard assays are available to study binding (e.g. surface plasmon resonance [31]) and catalysis (e.g. reaction product measurements in solution [32]), it is more challenging to produce a physiologically-relevant assay to explore the role of molecular reach. Recently, an *in vitro* reconstitution of the dephosphorylation of CD28 by PD-1 has been described whereby CD28 and PD-1 were localised to the two-dimensional surface of liposomes [7]. This system can be used to experimentally determine how changes to the molecular reach of the reaction influence the potency of PD-1. We have also recently introduced an SPR-based assay that can directly determine the molecular reach for fratricide reactions [9]. As these experimental tools mature, it may become feasible to systematically examine the role of molecular reach in controlling tethered signalling reactions.

## ACKNOWLEDGMENTS

SAI and YZ were supported by National Science Foundation award DMS-1255408. SAI was partially supported by a grant from the Simons Foundation, and thanks the Isaac Newton Institute of Mathematical Sciences for hosting him as a visiting Simons Fellow for the programme on Stochastic Dynamical Systems in Biology during which this project was initiated. OD is supported by a Wellcome Trust Senior Research Fellowship in Basic Biomedical Science (207537/Z/17/Z). JA is supported by a National Science Foundation grant DMS-1454739.

## SUPPORTING INFORMATION

### S1. Introduction to CRDME Model and Simulation Algorithm

The convergent reaction-diffusion master equation (CRDME) model we use corresponds to a spatial discretization of the general volume-reactivity model. In the latter, molecules are represented as point particles moving by Brownian motion [11, 33, 34]. First order reactions are modeled as internal processes with Poisson clocks, while bimolecular reactions between two molecules occur with a separation dependent probability per time (given by the Gaussian function *k*_cat_*σ*(*r*; *L*) for separation *r*, catalytic rate *k*_cat_, and molecular reach *L*, see Methods). These mathematical models can be described by their corresponding forward Kolmogorov equation, a high-dimensional coupled system of partial-integral differential equations for the probability density of having a given number of each chemical species at specified locations at a given time [11, 34]. For example, Eq. S14 is the forward Kolmogorov equation for the simplified case of just two molecules that can annihilate through the Gaussian interaction.

For multiparticle systems, the high-dimensionality of these equations precludes their solution by standard numerical methods for solving PDEs. Instead, we approximate the stochastic process of the individual molecules diffusing and reacting. In this work we do so by first spatially discretizing the forward Kolmogorov equation of the volume-reactivity model to a continuous-time Master Equation defined on a Cartesian mesh. We call this spatially discrete model the convergent reaction-diffusion master equation (CRDME) [10, 11]. As the set of ODEs that comprise the CRDME are still too high-dimensional to solve numerically, we instead generate exact realizations of the corresponding jump process associated with the CRDME using the Gibson-Bruck SSA method [35] (a variant of the well-known Gillespie method [36, 37]). We will subsequently call this simulation method the CRDME SSA. Here the diffusion of individual molecules is approximated by a continuous time random walk of the molecules hopping between voxels of the Cartesian mesh. Bimolecular reactions between reactant molecules in nearby voxels occur with probabilities per time derived from *σ*(·; *L*), see [10, 11] for full details.

As an illustrative example, consider a system with three chemical species, {A, B, C}, with each molecule diffusing within a square with periodic boundary conditions. Assume all molecules have diffusivity *D*, and the molecules may undergo the reactions 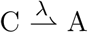 and 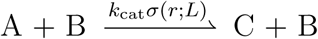 (where *r* denotes the separation of an individual pair of A and B molecules). To derive the CRDME model, we discretize the square into a collection of *N* square mesh voxels,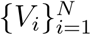, of width *h*. Let 𝒩(*V*_*i*_) label the set of the four nearest-neighbor voxels to voxel *V*_*i*_. The CRDME SSA then simulates the set of possible jump process transitions shown in Table S1. The bimolecular reaction transition rate in the table (i.e. probability per time) for one specific A molecule in *V*_*i*_ and one specific B molecule in *V*_*j*_ to react is given by

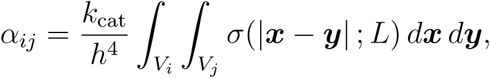

as derived in [11].

**TABLE S1.** Example of diffusive and chemical transitions in CRDME SSA simulations. For the 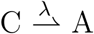 and 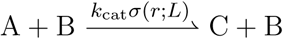 reaction-diffusion system, the table shows the five basic types of jump process transitions that can occur. Here *V*_*j*_ labels a given voxel of the Cartesian mesh, with mesh width *h* and four nearest-neighbors 𝒩 (*V*_*j*_). *A*_*j*_(*t*), *B*_*j*_(*t*) and *C*_*j*_(*t*) denote the stochastic processes for the number of molecules of each species in the *j*th voxel at time *t*. See SI S1 for details.

|  | Transitions | Transition Rates<br>(units of per time) | Upon Transition Event |
| --- | --- | --- | --- |
| Diffusive hopping<br>from $V_j$ to neighbor<br>$V_i \in \mathcal{N}(V_j)$: | $A_j \rightarrow A_i$ | $DA_j(t)/h^2$ | $A_i := A_i + 1, A_j := A_j - 1,$ |
| | $B_j \rightarrow B_i$ | $DB_j(t)/h^2$ | $B_i := B_i + 1, B_j := B_j - 1,$ |
| | $C_j \rightarrow C_i$ | $DC_j(t)/h^2$ | $C_i := C_i + 1, C_j := C_j - 1,$ |
| Chemical<br>Reactions: | $A_i + B_j \rightarrow C_i + B_j$ | $\alpha_{ij}A_i(t)B_j(t)$ | $A_i := A_i - 1, C_i := C_i + 1.$ |
| | $C_i \rightarrow A_i$ | $\lambda C_i(t)$ | $A_i := A_i + 1, C_i := C_i - 1.$ |

### S2. Derivation of the Doi model

We show here how to derive the simplified Doi model used in the Results section, beginning with a simplified model in which both the A and B molecules diffuse and interact through a Gaussian kernel. We again consider the two-particle annihilation reaction

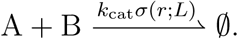

Assume the two molecules diffuse within a square (2.5D or 2D) or a cube (3D), Ω, where the length of each edge of the square (cube) is 300nm. Denote by *p*(***x, y***, *t*) the probability density an A molecule at ***x*** ∈ Ω and a B at ***y*** ∈ Ω have not yet reacted at time *t*. We consider the volume-reactivity model,

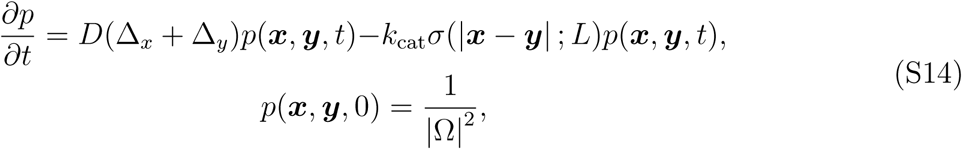

with reflecting boundary conditions on the boundary, *∂*Ω, in each of ***x*** and ***y***. Here *k*_cat_ is the catalytic rate of the reaction and *L* the molecular reach. |Ω| denotes the area (or volume) of Ω, so that the initial condition corresponds to starting both molecules well-mixed (i.e. uniformly distributed) within Ω.

We will focus on the behavior of the well-mixed mean reaction time (MRT), i.e. the average time for the diffusing A and B molecules to react assuming they are each placed randomly within Ω. Let *T* denote the random time at which the A and B molecules react. The exact well-mixed MRT is then given by the average of *T*, which we denote by ⟨*T*⟩. It is defined by

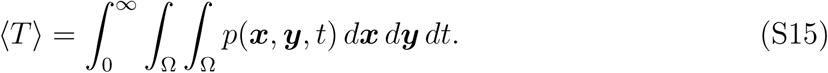

We simulate an approximation to the stochastic process of the two molecules diffusing and reacting using the CRDME SSA (see Methods and SI Section S1). Fig. S4 shows the estimated well-mixed MRT from 50000 CRDME SSA simulations in 2.5D. Observe that this figure is qualitatively quite similar to the asymptotic approximation to ⟨*T*⟩ found for the Doi model, Fig. 4B.

We now introduce two simplifications to the preceding model to obtain the Doi model. Due to presence of the 3D Gaussian interaction term *σ*(.; *L*) in Eq. S14, analytically solving for the exact ⟨*T*⟩ satisfying Eq. S15 is impractical. We therefore simplify Eq. S14 by transforming to a radially symmetric problem on a circle (sphere) of equivalent area (volume). In this reduced model one molecule is assumed stationary at the origin, while the other molecule diffuses with relative diffusivity *D* within a circle (sphere) of radius *R* about the origin. The MRT *u*(*r*) for a diffusing molecule that is initially placed a distance *r* from the origin satisfies

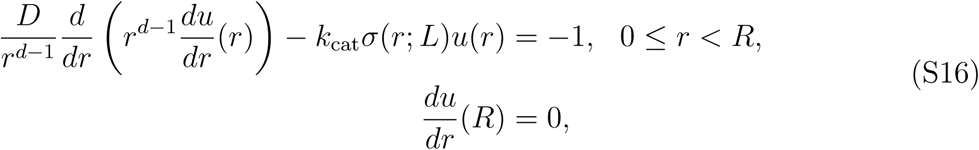

where *d* = 2 for the 2.5D and 2D models, and *d* = 3 for the 3D model. The corresponding well-mixed MRT when the position of the diffusing molecule is initially uniformly distributed is then given by

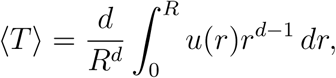

where again, *d* = 2 for the 2.5D and 2D models, and *d* = 3 for the 3D model. Eq. S16 is easily solved by a standard finite volume discretization, described in SI Section S3. Fig. S5 (solid lines) shows that in 2.5D this model gives almost identical results to the CRDME SSA approximation of Eq. S14 shown in Fig. S4.

To obtain an explicit analytical approximation to the solution of Eq. S16, we perform one final transform. We approximate Eq. S16 by a Doi model. As described in the Results section, the Doi model replaces the Gaussian interaction *σ*(*r*; *L*) by an approximating indicator function *λ*𝟙_[0,*ε*]_(*r*), giving the final model in Eq. 5. In the Results Section and SI Section S5 we describe how *λ* and *ε* are calibrated for all three cases. In the next section we show that the exact solutions in 2.5D, 2D and 3D agree well with the corresponding numerical solutions to the Gaussian interaction model (Eq. S16) over physiological parameter ranges.

### S3. Solving the radially (spherically) symmetric problem on a circle (sphere)

A key step in the analysis of the simplified model is to approximate the two-particle problem (Eq. S14) on a square (cube) with reflecting boundary conditions by a radially (spherically) symmetric problem (Eq. S16) with the same area (volume) and a reflecting boundary condition. Fig. S5A shows that the numerical solution of the PDE in Eq. S16 (solid lines) preserves the behavior of the 2.5D well-mixed MRT for the annihilation reaction obtained by CRDME SSA simulations of Eq. S14 (Fig. S4). We solved Eq. S16 numerically using a standard second-order finite volume discretization (in both the circle and sphere).

Similarly, in Eq. 5 we replace the Gaussian interaction of Eq. S16 by a calibrated Doi-model step-function interaction (see Eqs. 7 and S22). Fig. S5 shows that the exact solutions to the calibrated Doi model in Eq. 5 (dashed lines) in 2.5D (Fig. S5A), 2D (Fig. S5B), and 3D (Fig. S5C), each agree well with the numerical solution of the Gaussian interaction model (solid lines).

### S4. 3D well-mixed MRT for simplified Doi model

Using the same approach as for the 2.5D MRT analysis (see Results and S2), we map Eq. S16 to a 3D Doi type model with calibrated parameters that is analytically solvable. The 3D well-mixed MRT, ⟨*T*⟩, corresponding to substituting the solution of Eq. 5 into Eq. 8, is given by

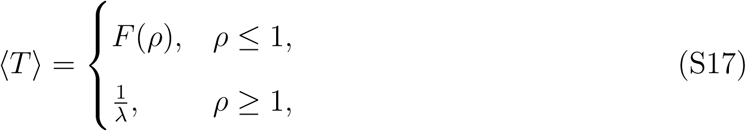

with 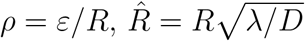 and

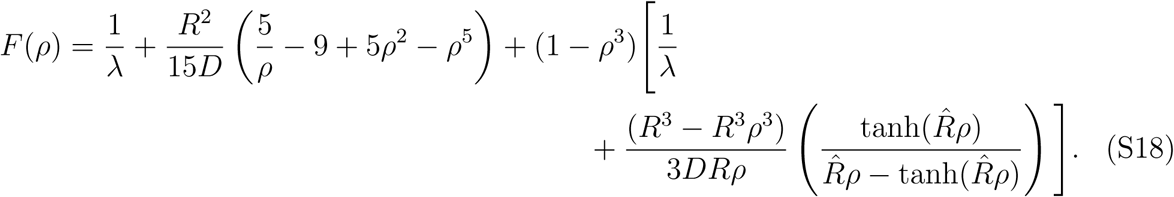

We use the same parameter calibration (Eq. 6) as we used for the 2.5D Doi model (Eq. 5) with 3D Gaussian interaction. *ε* and *λ* are then given by Eq. 7. Expanding Eq. S17 in *ρ* for *ρ* ≪ 1 we find

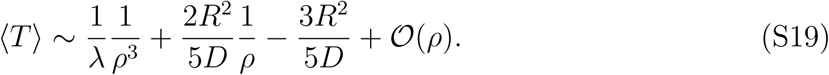

Substituting in the calibrated values for *λ* and *ε* then gives

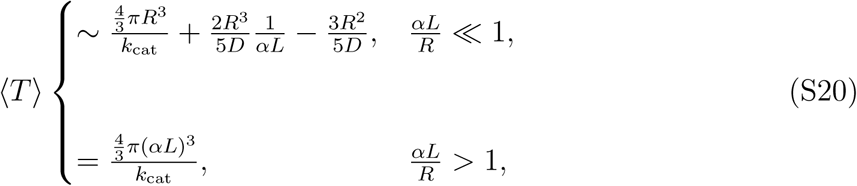

where 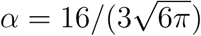.

### S5. 2D well-mixed MRT for a 2D concentration kernel

A key feature in all our models of tethered signalling is the use of a 3D Gaussian concentration kernel *σ*(*r*; *L*) to determine interaction functions for bimolecular reactions in the membrane. We now consider how the well-mixed MRT in the simplified 2.5D model (Eq. S16) changes if we instead use the 2D Gaussian interaction kernel *σ*_2D_(*r*; *L*) defined in Eq. 3. We call this new model the 2D model. We follow a similar analysis as in the preceding section and the Results.

The 2D well-mixed MRT, ⟨*T*⟩, obtained by substituting the solution of Eq. 5 into Eq. 8 is given by Eq. 9. As we now consider a 2D Gaussian interaction, we match the total area (equivalently total reaction rate) and the first moment of the 2D Gaussian using

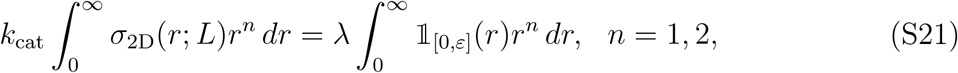

to obtain

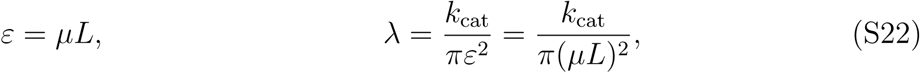

where 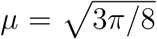. Notice, we now see the key difference from the use of a 3D interaction kernel; *λ* now scales like *L*^*−*2^ instead of *L*^*−*3^ as we previously found.

Expanding Eq. 9 in *ρ* for *ρ* ≪ 1 we again have Eq. 10, which combined with the preceding calibration for *ε* and *λ* then gives

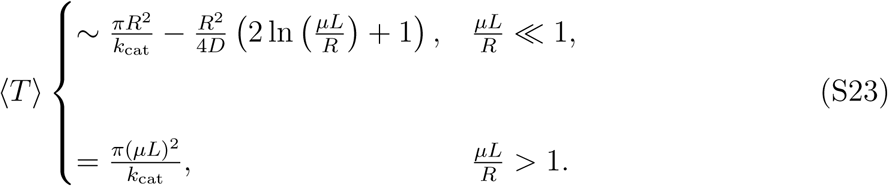

### S6. The well-mixed mean reaction time from the Doi model is approximately the sum of the reaction– and diffusion–limited mean reaction times

In each of the three asymptotic expansions Eqs. 12a, 12b and 12c, we now show the first two terms have a simple physical interpretation. The first term is essentially the mean reaction time if the system were reaction-limited (i.e. the well-mixed mean reaction time when *D* = *∞*). The second term is essentially the leading order diffusion-limited mean reaction time (i.e. the well-mixed mean reaction time when the reaction occurs instantly once the reactants are sufficiently close). We will illustrate how the former is responsible for the scaling regime where ⟨*T*⟩ grows in *L* for the (physiological) 2.5D model.

We first consider the well-mixed mean reaction time in the reaction limited regime. Assume that the diffusivity *D* of the diffusing molecule is infinite, so that the system is completely well-mixed. The position of the diffusing molecule is then given by a uniform density,

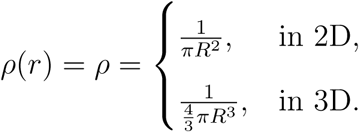

For each of the 2.5D, 2D and 3D models considered in the last Results section, in this regime we expect the total probability per time the molecules react to be given in terms of a well-mixed reaction-limited reaction rate, *k*_RL_, by

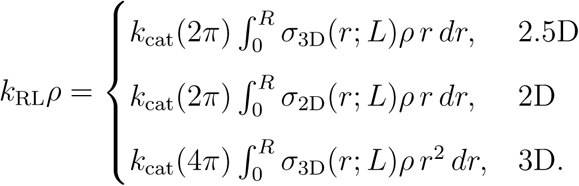

Assuming *R* is large, the integrals are well-approximated by the integral over *r* ∈ [0, ∞), so that

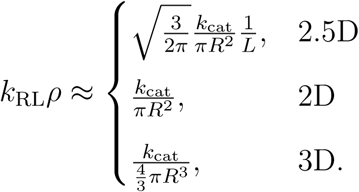

We note that the latter two are simply the standard probability per time a well-mixed reaction with bimolecular rate constant *k*_cat_ occurs within a circle (sphere) of radius *R*. The inverse of *k*_RL_*ρ* then defines the reaction-limited well-mixed mean reaction time,

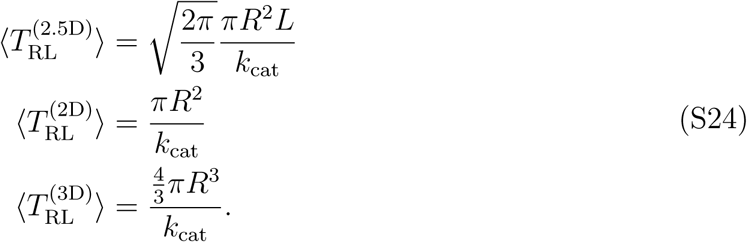

Only 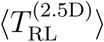 varies with *L*, increasing linearly as *L* increases.

Similarly, we may consider a diffusion limited regime in the (calibrated) Doi model, where the molecules react instantly upon the diffusing molecule reaching *r* = *ε*. The leading order asymptotic expansions for *ε/R* ≪ 1 of the diffusion limited well-mixed mean reaction time are well-known, see [38, 39], and given by

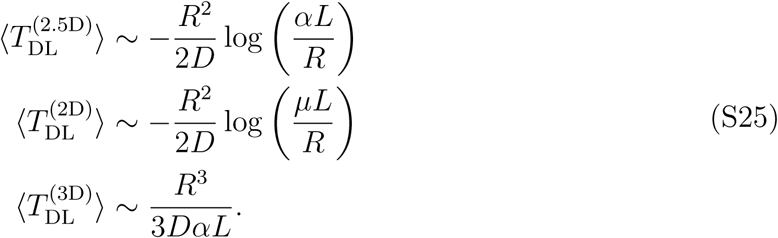

All three diffusion limited mean reaction-times are decreasing as *L* increases.

We therefore see that the first two terms in the asymptotic expansions of ⟨*T*⟩ from the Doi model (Eqs. 12a, 12b and 12c) can be summarized as essentially a sum of the reaction-limited and diffusion-limited well-mixed mean reaction times as given in Eq. 13.

### S7. Domain size dependence of the CD28 model

As a simple control, we solve the CD28 model (see Results) using a larger domain size to confirm that the steady-state fraction of phosphorylated CD28 exhibits a similar qualitative behavior with respect to the diffusivity and molecular reach. We see from Fig. S7C and S7D that solving the model on a larger square of side length of 500nm preserves the qualitative dependence of the steady-state fraction of phosphorylated CD28 on *D* and *L*.

### S8. Determining termination time in CRDME simulations

To determine the termination time in the CRDME SSA simulations of the first two models of the Results section, for each pair of diffusivity and molecular reach values we ran 100 test simulations. We then estimated an approximate time at which mean concentrations and/or fractions of each chemical species had reached steady state. In our final larger sampling runs (with *O*(10^4^) to *O*(10^5^) simulations per parameter set), we then set the termination time to be 40%-50% larger than the estimated time steady-state was reached. In Fig. S8 we plot till the termination time used in our final simulations the average fraction of phosphorylated CD28 in the CD28 model from 100 simulations (for different values of the diffusivity and molecular reach). In all nine cases the CD28 model appears to have reached steady state well before the chosen termination time.

### S9. Validity of using a stationary probability density kernel (*σ*)

We have used a stationary kernel (*σ*) to represent the probability that two tethers interact. This assumption is only valid when the timescale over which the polymer tether explores its space (*τ*) is smaller than the timescale for reaction and diffusion. An approximate upper bound on *τ* can be obtained by assuming the polymer is a freely-jointed chain with *N* Kuhn segments of length *b*, and monomer size *< b*. These are related to the reach parameter, *L*, by 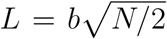. A lower bound on the diffusion coefficient of the polymer is 6*πηL/k*_*B*_*T*, leading to the following upper bound estimate,

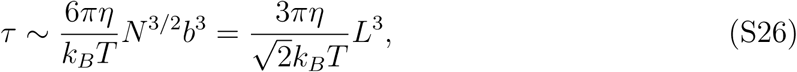

where *η* is the viscosity of the surrounding medium. Assuming that the viscosity of the cytoplasm is close to that of water, we find that *τ* ≲ 10^*−*4^ s for the maximum molecular reach of *L* = 32 nm in Fig. 2. This is smaller than the fastest diffusive timescale (*∼* L^2^/D = (32 nm)^2^/0.125 *µ*m^2^/s = 8 *×* 10^*−*3^ s) and reaction timescale 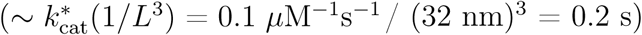.

The above approximation has been derived in a more general setting [40, 41] and confirmed in numerical simulation [41–43]. This model was quantitatively confirmed and in particular the diffusion coefficient equation [44] and the loop closure time equation [45] were found to be in close agreement. The scaling law *τ ∼ N* ^3*/*2^ was confirmed experimentally [46] and by all-atom dynamics [47].

### S10. Reaction kernel for surface-tethered molecules

As discussed in the Materials & Methods, if two tethers have their bases fixed at a separation distance *r*, then the probability density for the reaction sites of the tethers to interact is given by Eq. 1, which is referred to as the reaction kernel. This equation was derived by assuming each tether *i* explores three-dimensional space with Gaussian probability density. We note that this formulation assumed that the tether was allowed to explore all of space. In this section, we ask, what is the interaction kernel *σ*_surf_ (*r*) for two tethers with base fixed at a distance *r*, but which are attached to a 2D surface (i.e. the plasma membrane), and therefore can only explore half-space.

Since surface-adhered polymers do not obey Gaussian probability densities that only depend on *L*, we must specify more detailed polymer properties. We assume the tethers are freely-jointed chains composed of *N* rigid segments of length *δ*. In free-space, this chain has a reach parameter of 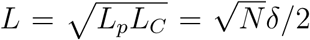. We assume *δ* = 0.3 nm consistent with previous models of disordered proteins and PEG [48, 49]. We explore a range of tether lengths *N* = 25 *−* 1000 corresponding to free-space reach parameters *L* = 1.5 *-* 13 nm. We simulate the ensemble of polymer conformations of the two freely-jointed chains, for various base separation distances *r*, using a standard Metropolis method [43, 50] and determine the probability that the reaction sites interact to determine *σ*_surf_.

In Fig. S9A we show the probability density of the end-to-end distances for various values of *L*. In all cases, the distances are slightly elongated by the presence of the surface. This is in agreement with previous findings that adherence to a surface tends to elongate polymers [51]. In Fig. S9B, we show the reaction kernel *σ*_surf_ (*r*). In all cases, we find that the effective concentrations are enhanced by the presence of the surface. This makes intuitive sense, since reaction sites are forced by the surface to be in the same half-space. The length-scale of the separation distance, i.e., the decay length of the curves in Fig. S9B, is approximately unchanged by the surface. The simulation therefore suggests that the reaction kernel for surface-bound tethers is well-approximated by

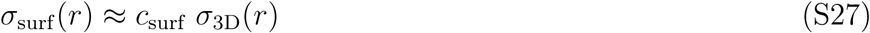

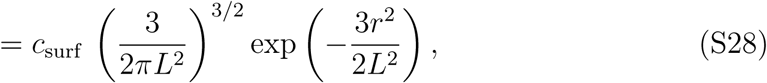

where *c*_surf_ *>* 1 is an enhancement factor that arises as the surface forces the tethers together.

By fitting Eq. S28 to the simulated kernels in Fig. S9B, we produce estimates for the enhancement factor *c*_surf_ in Fig. S9C. We find that *c*_surf_ *≈* 1.5 over the estimated physiological range of *L* (for 7 nm*< L <* 13 nm, always within 10%). In CRDME simulations of reactions between tethered molecules, the per-second reaction rate is *k*_cat_*σ*_3D_(*r*). The results here allow us to use the same scheme to simulate reactions between surface-tethered molecules, but with the prefactor *k*_cat_ reduced by *≈* 1.5.

To further verify the validity of this approximation, in Fig. S9D we plot the re-scaled kernel

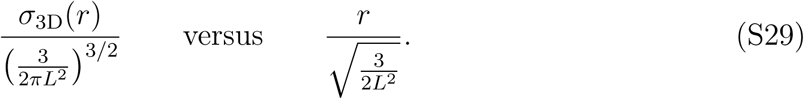

In this re-scaling, all free-space kernels collapse onto a single curve (black curve in Fig. S9D). We now plot the surface-adhered kernels re-scaled as

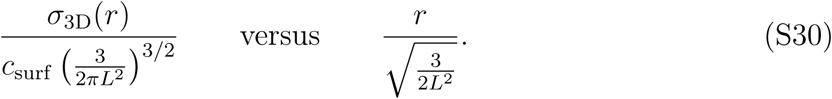

We find that these surface-adhered kernels also collapse onto the same single curve. This confirms the approximation given by Eq. S28.

---

**FIG S1.**
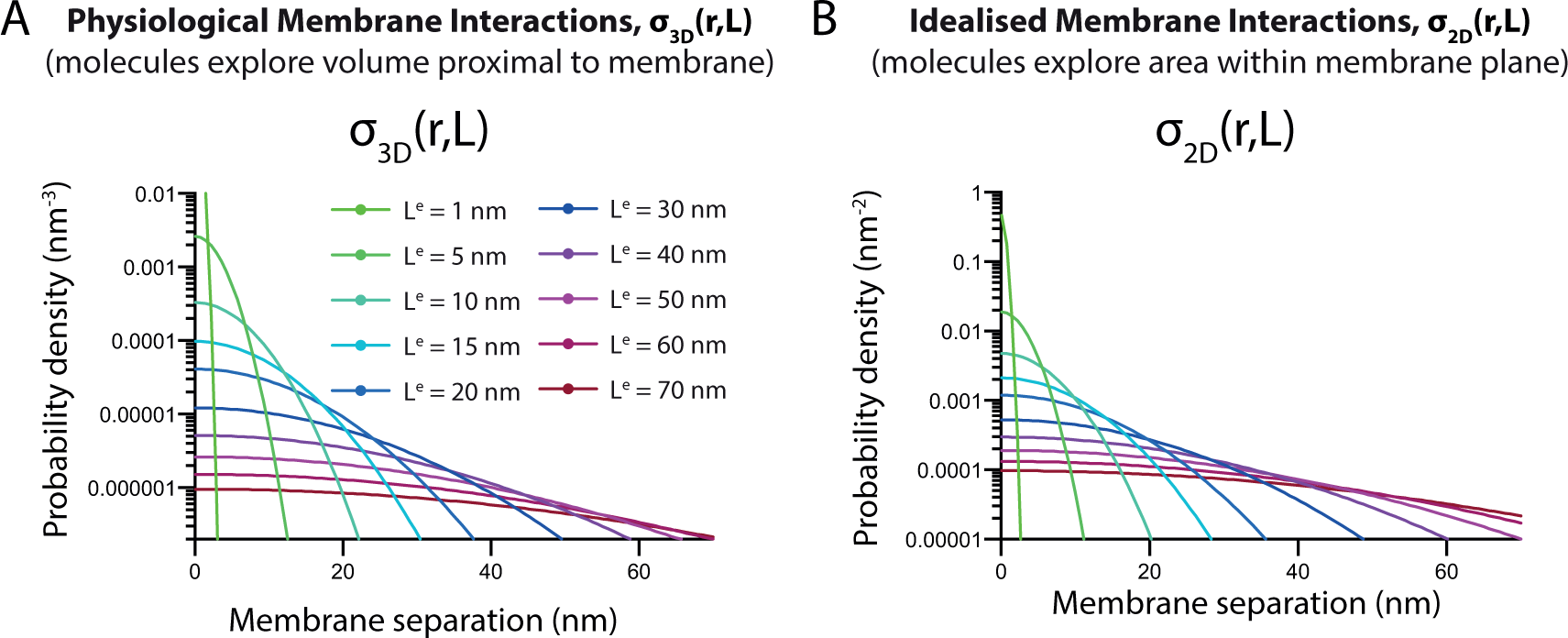
Comparison of A) *σ*_3D_ and B) *σ*_2D_ over the membrane separation distance (nm) for the indicated value of the molecular reach of the reaction (*L*).

**FIG S2.**
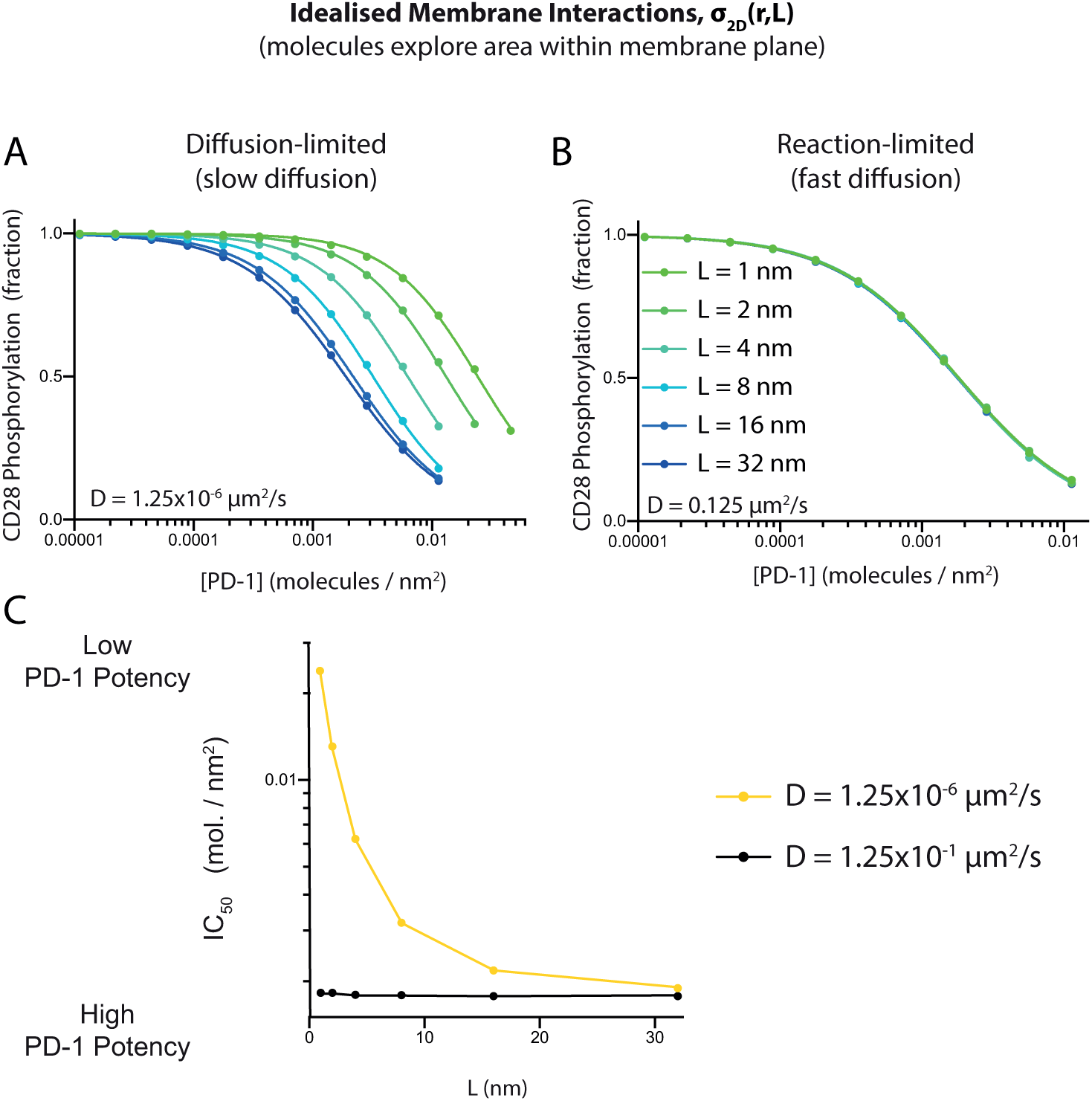
Reproducing Fig. 2 using the idealised 2D interaction kernel *σ*_2D_ shows that increasing the molecular reach A) increases PD-1 potency in the diffusion-limited regime but B) has no effect in the reaction-limited regime. C) The potency over the molecular reach quantified from A and B.

**FIG S3.**
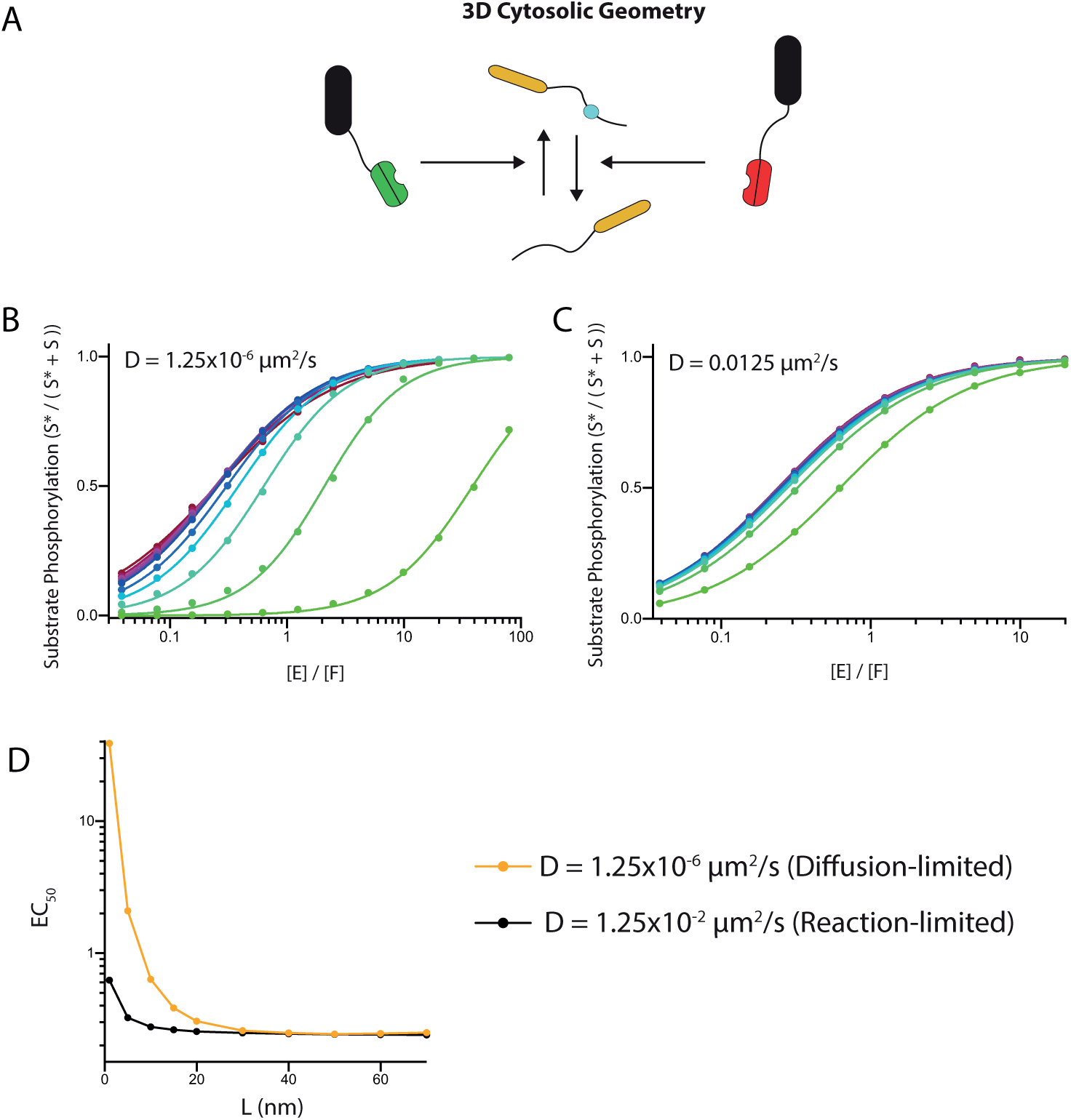
Reproducing Fig. 3A,C,D,E when molecules freely diffuse in 3D using the physiological 3D interaction kernel *σ*_3D_. A) Schematic of model highlighting that molecules are not confined to a membrane. B-C) Fraction of phosphorylated substrate in the steady-state for the indicated values of the molecular reach of the reaction when reactions B) are limited by diffusion or C) are not limited by diffusion. D) The ratio of kinase-to-phosphatase that produces half-maximal phosphorylation over the molecular reach of the reaction showing that increasing the molecular reach can only increase potency in this geometry. Parameter values: [S] = 8.5 *×*10^4^*µ*m^*−*3^, [F] = 9.4*×*10^4^*µ*m^*−*3^, domain size = 300nm *×* 300nm *×* 300nm, and all other parameters as indicated in Table II.

**FIG S4.**
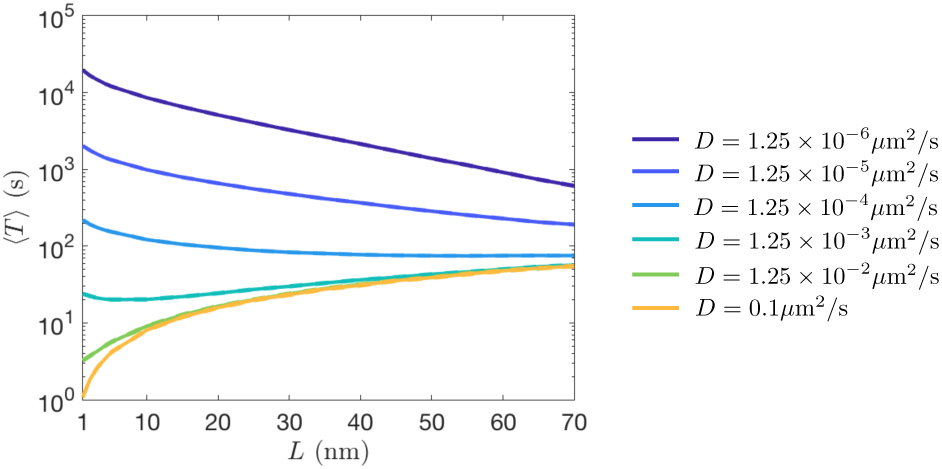
The well-mixed mean reaction time, ⟨*T* ⟩, of the two-particle 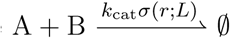 reaction changes its dependence on molecular reach for small vs. large diffusivities when the molecules diffuse in a (2D) membrane, but their tails can react in the (3D) cytosol. For each value of *D* and *L* we estimated ⟨*T* ⟩ from 50000 CRDME-SSA simulations. 95% confidence intervals for each curve are given by dashed lines of the same color (barely visible). The catalytic rate *k*_cat_ was 0.1 *µ*M^*−*1^s^*−*1^.

**FIG S5.**
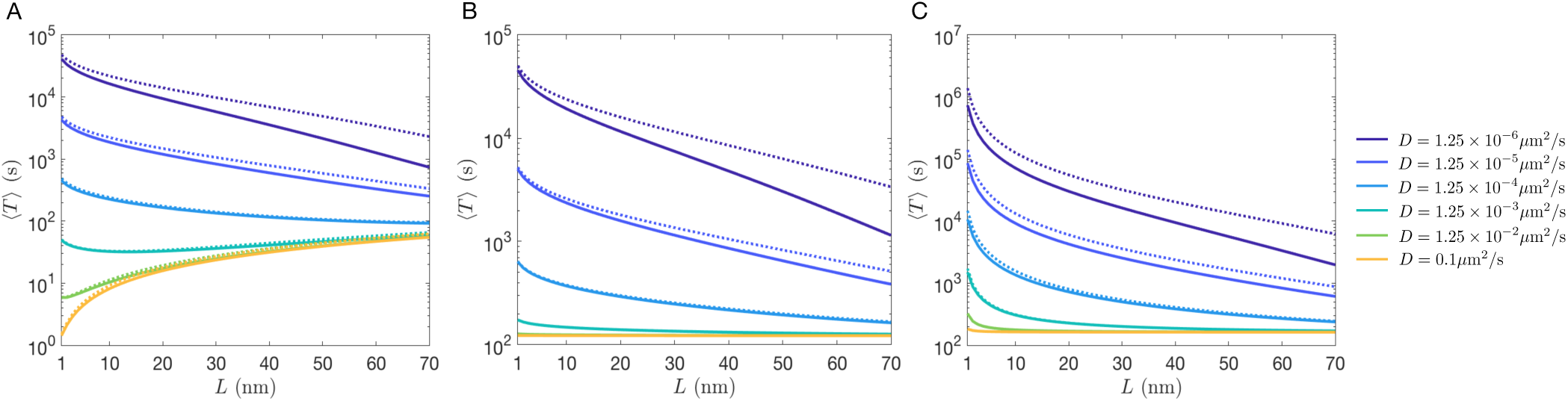
The well-mixed MRT ⟨*T* ⟩ determined from the numerical solution of the Gaussian interaction model given by Eq. S16 (solid lines) and exact solution to the Doi step function interaction model given by Eq. 5 using calibrated *λ* and *ε* values (dashed lines). See discussion of SI Section S3.(A) 2.5D model, having Doi solution (Eq. 9) and calibration (Eq. 7); (B) 2D model, having Doi solution (Eq. 9) and calibration (Eq. S22); (C) 3D model, having Doi solution (Eq. S17) and calibration (Eq. 7). The area (2D)/volume (3D) of the circle/sphere is chosen to be the same as the square/cube of side length 300 nm. For A and C the catalytic rate *k*_cat_ is set to be 0.1 *µ*M^*−*1^s^*−*1^. For B the catalytic rate *k*_cat_ is 0.1*πR*^2^*/*(4*πR*^3^*/*3) *µ*M^*−*1^s^*−*1^nm^*−*1^, the corresponding 2D bimolecular rate constant to A and C.

**FIG S6.**
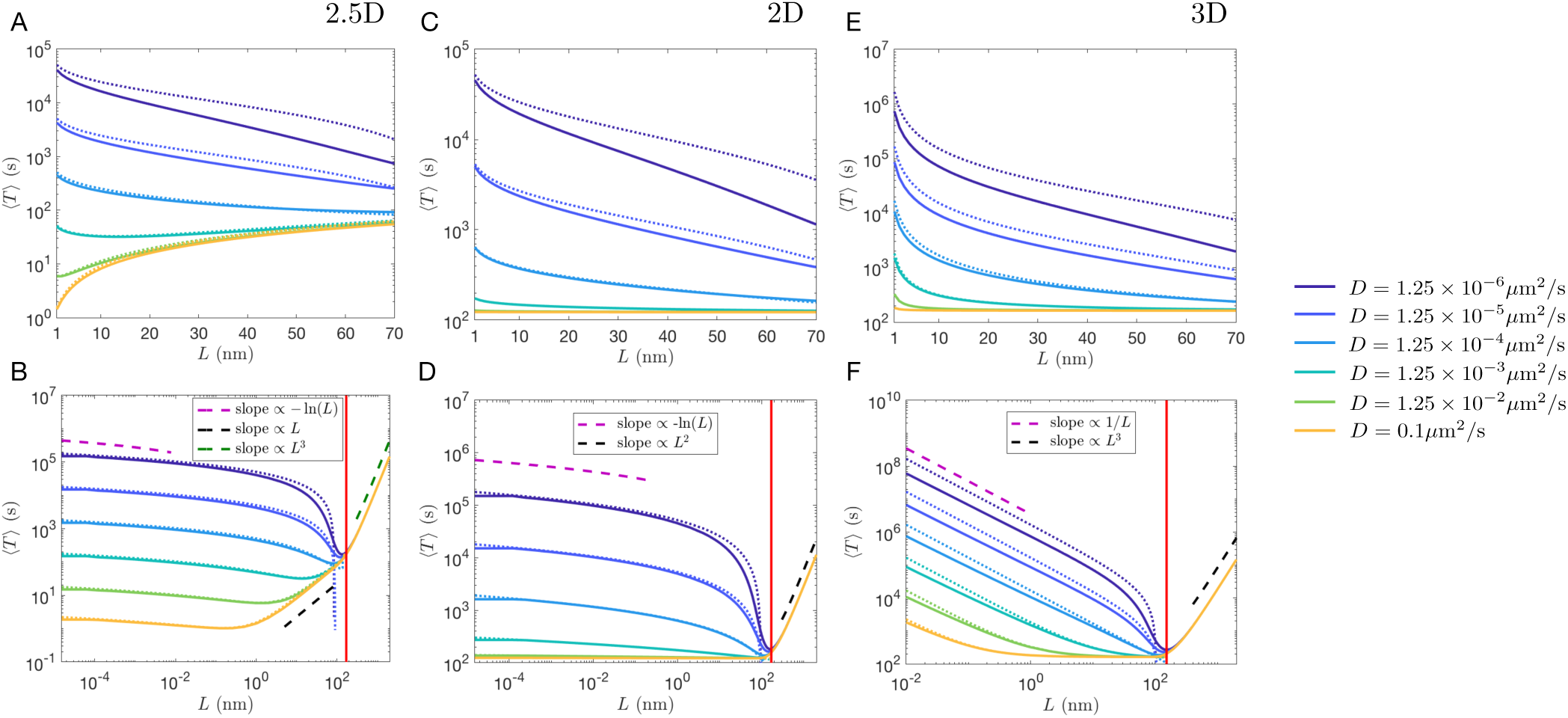
The well-mixed mean reaction time (MRT), ⟨*T*⟩, only demonstrates a switch in dependence on molecular reach for small vs. large diffusivities when considering membrane-bound molecules with cytosolic tails that react in 3D (2.5D model). In all figures solid lines correspond to ⟨*T*⟩ as estimated by numerically solving the ODE in Eq. S16. Dotted lines correspond to the asymptotic expansions in Eq. 12a for A/B, Eq. 12b for C/D and Eq. 12c for E/F. Dashed lines show general scaling behavior as a function of *L*. A) 2.5D model well-mixed MRT over physical parameter range. B) Same as A but showing an expanded range of *L* values. C) 2D model well-mixed MRT over physical parameter range. D) Same as C but showing an expanded range of *L* values. E) 3D model well-mixed MRT over physical parameter range. F) Same as E but showing an expanded range of *L* values. In the expanded range figures the red solid line gives the *L* value such that *ε/R* = 1, corresponding to when the Doi interaction distance, *ε*, is equal to the domain radius, *R*. Note, as *ε → R* from below the asymptotic expansions break down as 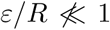. For A, B, E, and F the catalytic rate *k*_cat_ is 0.1 *µ*M^*−*1^s^*−*1^. For C and D the catalytic rate *k*_cat_ is 0.1*πR*^2^*/*(4*πR*^3^*/*3) *µ*M^*−*1^s^*−*1^nm^*−*1^, the analogous 2D catalytic rate.

**FIG S7.**
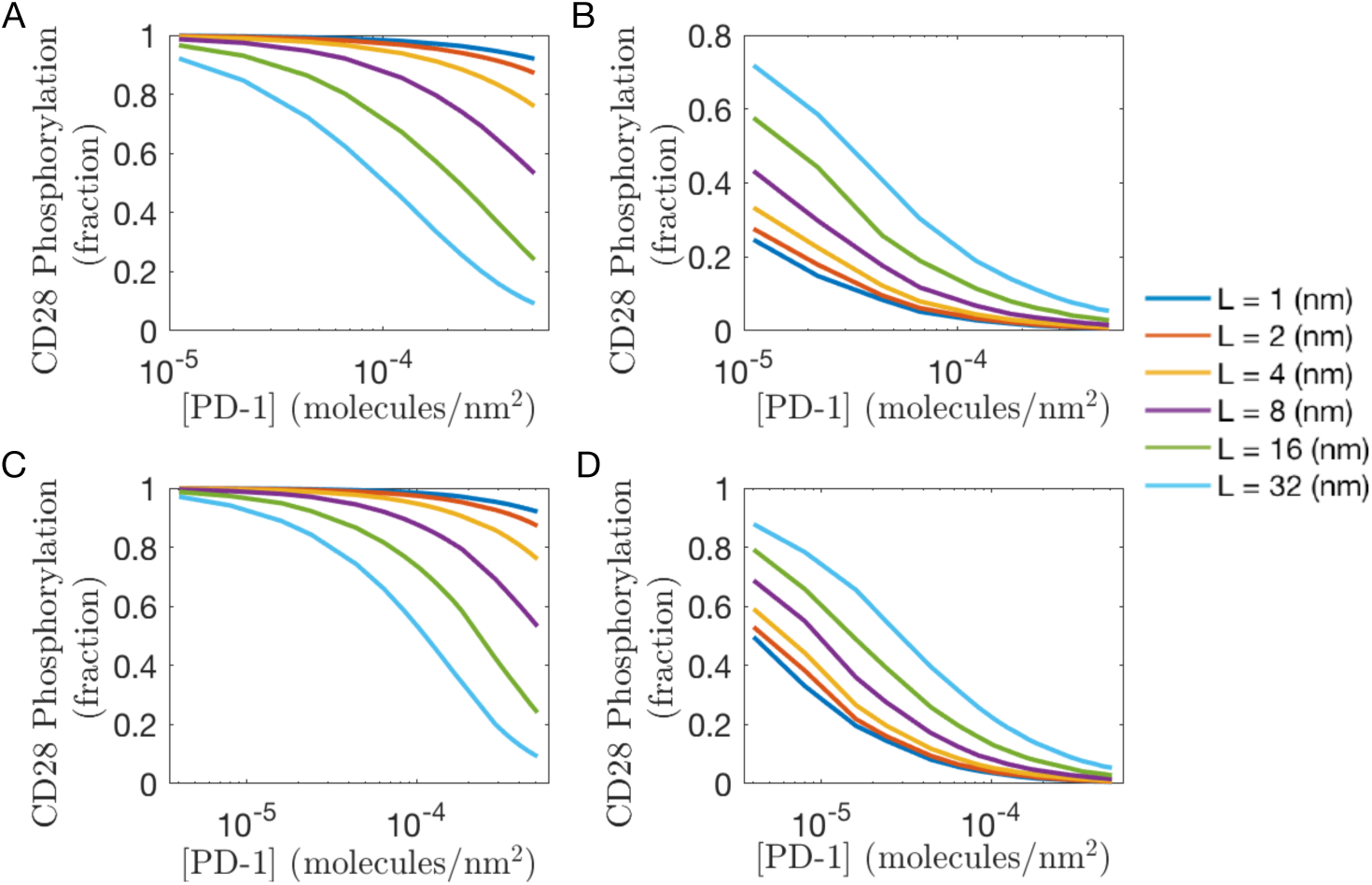
Dependence of the steady-state fraction of phosphorylated CD28 in the CD28 model for two different domain sizes when diffusion is slow (A,C; *D* = 1.25 *×* 10^*−*6^*µm*^2^*/s*) and fast (B,D; *D* = 1.25 *×* 10^*−*2^*µm*^2^*/s*). A) and B) The steady-state fraction of phosphorylated CD28 using a square domain of side length 300nm. C) and D) The steady-state fraction of phosphorylated CD28 using a square domain of side length 500nm. [PD-1] is shown on a logarithmic scale in each figure. Each curve was estimated from 100000 simulations. The catalytic rate 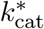 was set to be 0.01*µ*M^*−*1^s^*−*1^.

**FIG S8.**
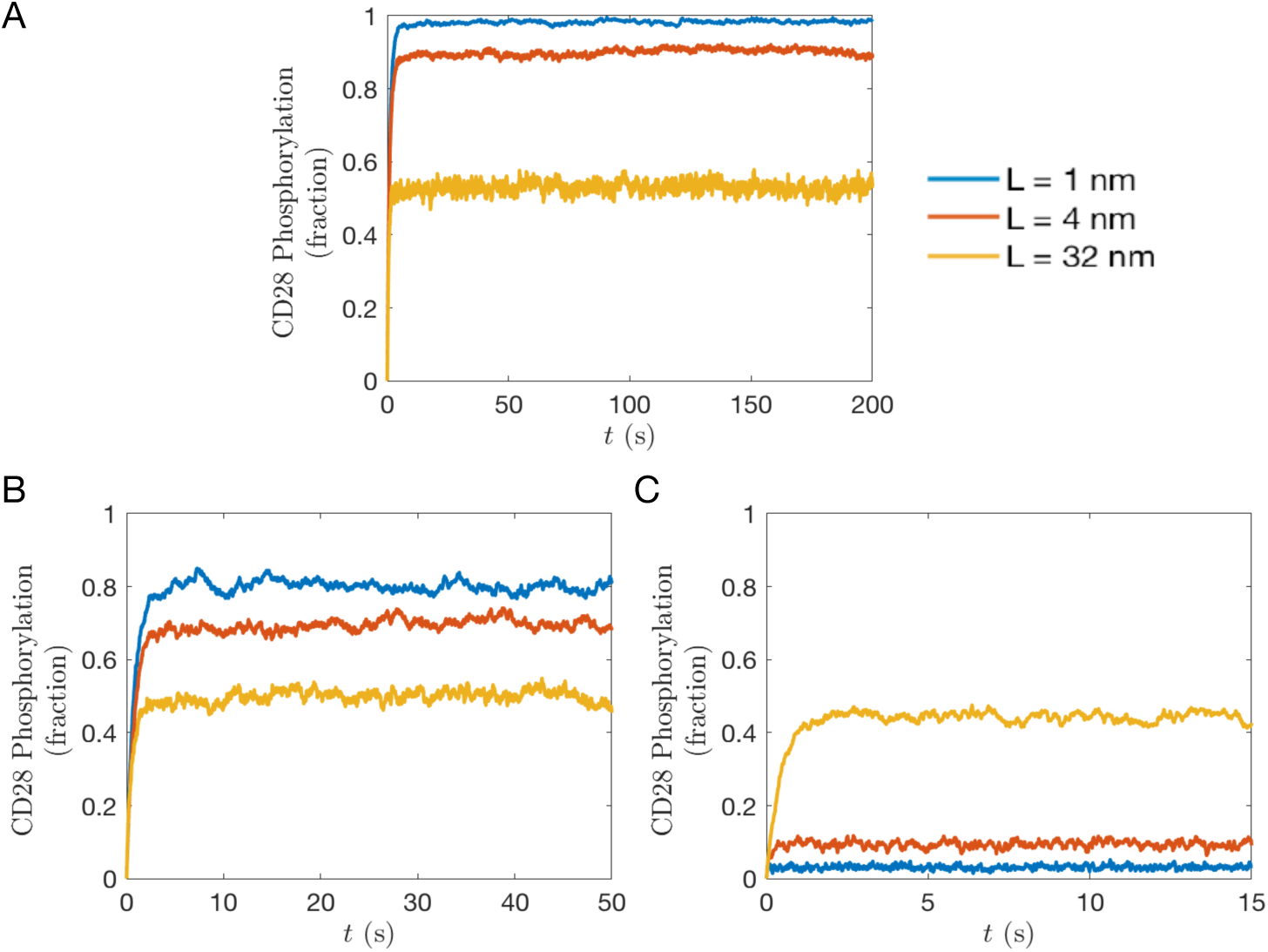
Average fractions of phosphorylated CD28 in the CD28 model versus time. Each figure ends at the termination time at which we concluded the system had reached steady-state. These times were then used in larger sets of simulations to produce Fig. 2. A) *D* = 1.25 *×* 10^*−*6^*µ*m^2^*/s*. B) *D* = 1.25 *×* 10^*−*4^*µ*m^2^*/s*. C) *D* = 1.25 *×* 10^*−*1^*µ*m^2^*/s*. Each curve is an average from 100 simulations with an initial PD-1 concentration of 3.5556 *×* 10^*−*4^*/*nm^2^. Remaining parameters are as in Table I.

**FIG S9.**
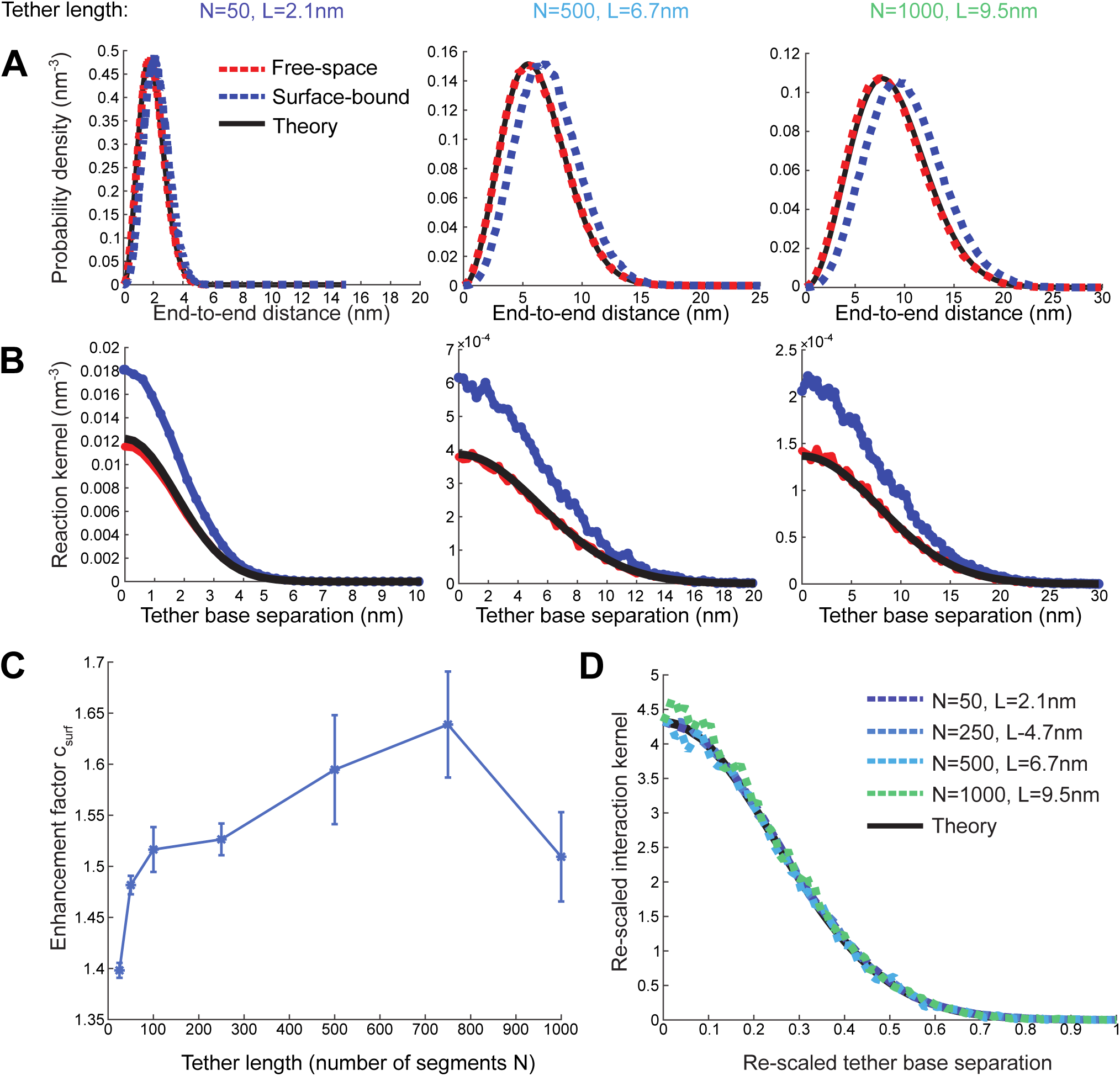
Simulation of tethers of various lengths adhered to a surface. (A) End-to-end distances for surface-bound tethers (blue dashed) have probability densities shifted slightly larger than free-space (red dashed). Free-space results can be compared with analytical theory (black solid) for validation. Reaction kernel for surface-adhered tethers *σ*_surf_ (*r*) (blue-dashed) and free-space *σ*_3D_(*r*) (red dashed). The surface enhances the reaction kernel by approximately 1.5-fold. Free-space result is compared to analytical theory (black solid). (C) Best-fit factor *c*_surf_ that fits the surface-adhered kernel (blue curves in (B)) to the approximation (Eq. S28). (D) If the reaction kernels from (B) are re-scaled according to Eq. S30, we find that all kernels, for the full range of tether length *N* we explored, collapse onto a single curve, demonstrating the validity of the approximation (Eq. S28) with this choice of enhancement factor *c*_surf_.

